# Patterns of thermal adaptation in a worldwide plant pathogen: local diversity and plasticity reveal two-tier dynamics

**DOI:** 10.1101/2019.12.16.877696

**Authors:** Anne-Lise Boixel, Michaël Chelle, Frédéric Suffert

## Abstract

- Plant pathogen populations inhabit patchy environments with contrasting, variable thermal conditions. We investigated the diversity of thermal responses in populations sampled over contrasting spatiotemporal scales, to improve our understanding of their dynamics of adaptation to local conditions.
- Samples of natural populations of the wheat pathogen *Zymoseptoria tritici* were collected from sites within the Euro-Mediterranean region subject to a broad range of environmental conditions. We tested for local adaptation, by accounting for the diversity of responses at the individual and population levels on the basis of key thermal performance curve parameters and ‘thermotype’ (groups of individuals with similar thermal responses) composition.
- The characterisation of phenotypic responses and genotypic structure revealed: (i) a high degree of individual plasticity and variation in sensitivity to temperature conditions across spatiotemporal scales and populations; (ii) geographic adaptation to local mean temperature conditions, with major alterations due to seasonal patterns over the wheat-growing season.
- The seasonal shifts in functional composition suggest that populations are locally structured by selection, contributing to shape adaptation patterns. Further studies combining selection experiments and modelling are required to determine how functional group selection drives population dynamics and adaptive potential in response to thermal heterogeneity.

## Introduction

Populations in natural settings often experience environmental heterogeneity (Li & Reynolds, 1995), which dictates physiological responses (Cavieres & Sabat, 2008) and can drive the emergence of local adaptation patterns (Thompson, 2005; Nuismer & Gandon, 2008). As such, heterogeneity is regarded as one of the most important elements driving the emergence and maintenance of genetic variation within populations (Levins, 1974; Hedrick, 1986; Ravigné *et al*., 2009), thereby shaping population dynamics (Hughes *et al*., 2008). Gathering information about the way a given community, species or population copes with this environmental heterogeneity is crucial for the understanding and prediction of its distribution and responses to current and future environmental changes (Austin, 2007).

The adequate capture of eco-evolutionary responses requires an integration of physiological variation across biological (individual, group, population, species) and spatiotemporal (seasonal, geographic) scales, given the significant implications of this variation for dynamics (Saloniemi, 1993; Vindenes *et al*., 2008; Schreiber *et al*., 2011). It is therefore important to go beyond summarising diversity through average trait values (Bolnick *et al*., 2011; Violle *et al*., 2012), and to account for the individual specialisation of phenotypic responses by taking into account both phenotypic plasticity (within-individual differences; Pigliucci, 2001) and interindividual variation (between-individual differences; Dall *et al*., 2012). This paradigm shift has been made possible by progress in the measurement and analysis of this specialisation at the individual and population level.

The ecological concept of “reaction norm”, describing the set of phenotypes generated by a given genotype in response to environmental cues (Schlichting & Pigliucci, 1998), is particularly effective for accounting for individual specialisation (Bolnick *et al*., 2002; Araújo *et al*., 2008). Most of the comparisons on the continuous variation of a given phenotypic trait among individuals under different environmental conditions have been conducted to date on reaction norm descriptors (*e.g.* comparisons of phenotypic mean and range values at the maximum; Gibert *et al*., 1998) or degree of plasticity (*e.g.* breakdown of the shifting and stretching of non-linear reaction norms into non-exhaustive biological modes, *i.e.* lower-higher, faster-slower, specialist-generalist directions; Izem & Kingsolver, 2005; Martin *et al*., 2011; van de Pol, 2012). Such approaches have proved highly valuable, but may not be suitable for decomposing the overall variation or distinguishing differential responses between populations (Bulté & Blouin-Demers, 2006) or including intra- and interindividual sources of error (Angilletta, 2006) in ANOVA and random regression approaches (Lynch & Gabriel, 1987; Gilchrist, 1995).

One possible complementary approach to the description of variation between reaction norms involves the use of functional ecology to describe significant variations in the intensity of individual specialisation within populations and species (Garnier & Navas, 2012). The idea is to translate reaction norms into functional traits (Violle *et al*., 2007) by grouping individual reaction norms into ‘functional groups’ (Gitay & Noble, 1997). Each of these functional groups responds to the environment in its own way (*e.g.* low- or high-performance specialists), according to a classification system that is not predetermined (*i.e.* constrained modes of variation). This approach accounts more effectively for diversity and adaptation patterns, through the characterisation of three functional components: richness, evenness, and divergence (Mason *et al*., 2005).

This approach is particularly useful for deciphering variation in continuous reaction norms describing performance as a function of temperature (thermal performance curves or TPC; Huey & Stevenson, 1979), and for documenting patterns of thermal adaptation to prevailing local conditions (Kawecki & Ebert, 2004) across a range of environments (*e.g.* Mitchell & Lampert, 2000). These patterns plays an important role in the case of microorganisms impacting ecosystems, human health, and food security (Fisher *et al*., 2012) as local adaptation to temperature conditions governs their geographic distribution, phenology, and abundance (Kraemer & Boynton, 2017). This results in impacting the expansion ranges of plant pathogens (*e.g.* Milus *et al*., 2009; Robin *et al*., 2017), as well as the onset and severity of disease epidemics (*e.g.* Ferrandino, 2012).

The studies of thermal responses in plant pathogenic microorganisms performed to date have focused mostly on either summarising the individual variance of aggressiveness traits as population-scale averages (problematic use of single mean species values; Suffert & Thompson, 2018) or phenotyping individuals under a limited set of temperatures when considering variances (generally about three temperatures in thermal biology studies; Dell *et al*., 2013; Low-Décarie *et al*., 2017). These strategies have provided useful information about species distribution, making it possible to detect signatures of interindividual variation and adaptation within species and populations (Milus *et al*., 2006). However, they cannot be used to infer group selection driving population dynamics (Lavergne *et al*., 2010) or to assess the relevant scales of functional diversity (Woodcock *et al*., 2006; Martiny *et al*., 2011). Such analyses go well beyond simple comparisons of mean trait values and would require the characterisation of entire TPCs and their variation across different scales.

This study develops a functional approach ascertaining patterns of diversity across space (geographic range) and time (local seasonal dynamics) in a way that explores how functional diversity in thermal responses varies with environmental conditions and uncovers the role that adaptation plays in generating this diversity. The analysis of the plasticity and variation of thermal sensitivity across individuals, populations, and scales was conducted in the case of the fungal wheat pathogen *Zymoseptoria tritici* (formerly *Mycosphaerella graminicola*; Steinberg, 2015). This pathogen, the causal agent of one of the most economically important wheat diseases (Septoria tritici blotch or STB; Dean *et al*., 2012; Fones & Gurr, 2015), is now considered a model species for both basic and applied research. Besides its agronomic relevance, we chose to study *Z. tritici* as its aggressiveness traits are known to be temperature-sensitive (Shaw, 1990; Lovell *et al*., 2004) and to display interindividual variation (Bernard *et al*., 2013; Boixel *et al*., 2019). Furthermore, its populations present signatures of adaptation to a wide range of contrasted environments over space (wheat-growing areas worldwide; Zhan & McDonald, 2011) and time (covering seasonal changes *e.g.* from late autumn to early summer in Europe; Suffert *et al*., 2015). Drawing on previous local adaptation studies conducted by Zhan & McDonald (2011) and Suffert *et al*. (2015), we designed a sampling scheme to grasp the levels of functional diversity shaping responses of its populations to contrasted environments based on a finer-grained sampling and phenotyping resolution.

## Materials and Methods

### Tailored sampling survey design (Fig. 1 – step 1)

Samples were collected from 12 *Z. tritici* populations for the exploration of spatial and temporal components of thermal adaptation. Spatial variation was investigated within the Euro-Mediterranean region, with samples from eight sites in wheat-growing areas representative of the highly contrasting climatic conditions over this large geographic area (see the 8 populations for the geographic scale, covering three out of seven Köppen-Geiger climate zones in which *Z. tritici* is reported as a notable pathogen, in Table 1 and Fig. S1). One of these sites (Grignon, France) was selected for a comparison of the thermal responses of two pairs of winter and spring subpopulations sampled from neighbouring fields, to capture seasonal dynamics over a wheat growing season (*i.e.* over the course of an annual epidemic; see the 4 populations of the seasonal scale in Table 1 and Fig. S2). These pairs of subpopulations were subject to seasonal variation from November to February and from March to June, respectively. For each of the 12 populations, we collected from 25 to 30 isolates at random from wheat leaves with STB symptoms, from which single-spore isolates were prepared (Methods S1) and which were later confirmed to be genetically unique strains with the microsatellite analysis. We chose to consider 25 or 30 strains (*i.e.* individuals) per population instead of the minimum level of 15 identified on the basis of a rarefaction analysis (Fig. S3) for estimating the diversity of thermal responses in *Z. tritici* with sufficient power, accuracy and precision (Dale & Fortin, 2014).

**Fig. 1.**
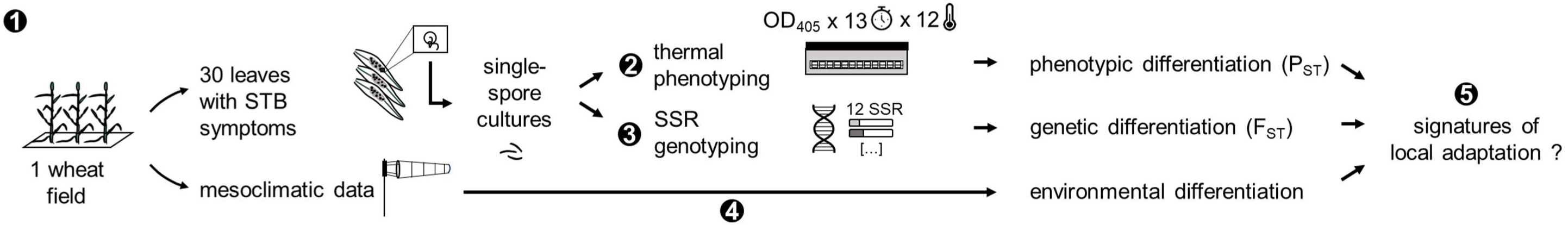
Overview of the methodology for characterising diversity and adaptive patterns of thermal responses in the sampled *Zymoseptoria tritici* populations. (Step 1) 12 populations, each composed of either 25 or 30 strains, were collected from diseased leaves in different spatiotemporal locations (8 Euro-Mediterranean populations collected along geographic thermal gradients and 4 French seasonal subpopulations) with the corresponding mesoclimatic conditions (temperature data). All strains were: (Step 2) phenotyped in an *in vitro* growth experiment conducted over a range of 12 temperatures to capture thermal performance curves from growth kinetics involving 13 measurement time points (experimental framework detailed in Boixel *et al*., 2019); (Step 3) genotyped for 12 neutral microsatellite (SSR) markers to quantify phenotypic (P_ST_) and genetic (F_ST_) differentiation. (Step 4) The thermal conditions experienced by individuals over the wheat growing season were characterised for each spatiotemporal site. (Step 5) The local adaptation of individuals and populations to temperature was assessed by cross comparisons of the spatiotemporal patterns of thermal responses, allele frequency and thermal conditions.

**Table 1.**
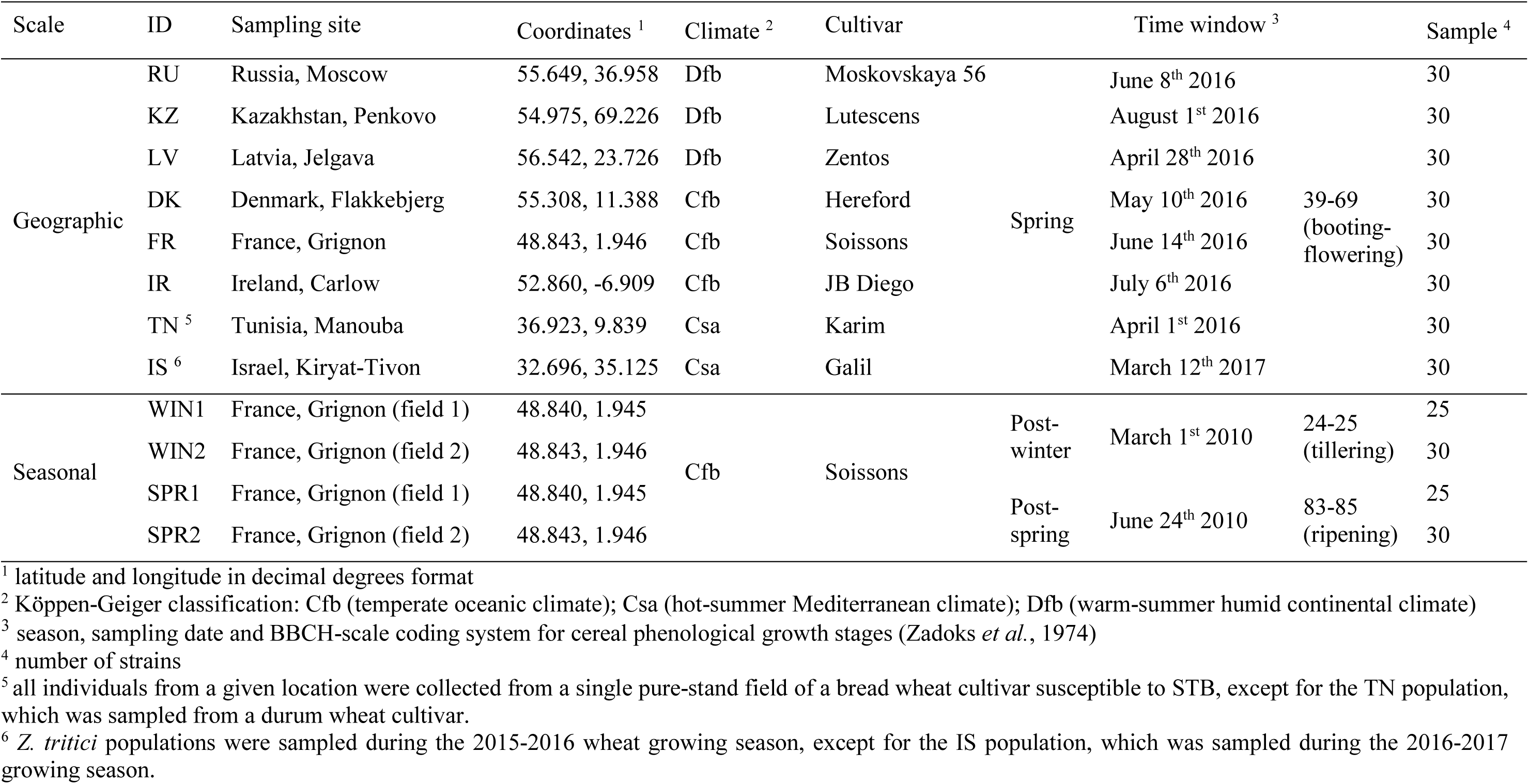
Summary information for the 12 Zymoseptoria tritici populations. These populations were specifically sampled along geographic (8 Euro-Mediterranean populations) and seasonal (4 local French winter (WIN) and spring (SPR) subpopulations) scales with contrasting temperature conditions. Strains were collected from naturally infected wheat fields characterised by a spatial location, a climate zone, a wheat cultivar and sampling conditions (time window and sample size).

### Phenotypic variations in thermal responses (Fig. 1 – step 2)

Thermal responses were phenotyped by determining the *in vitro* growth rates of the strains in liquid glucose peptone medium (14.3 g.L^−1^ dextrose, 7.1 g.L^−1^ bactopeptone and 1.4 g.L^−1^ yeast extract) over a four-day period at 12 temperatures ranging from 6.5 to 33.5°C (6.5, 9.5, 11.5, 14.5, 17.5, 20.0, 22.5, 24.5, 26.5, 28.5, 30.5 and 33.5°C). The growth rate *μ* of each strain at each temperature was calculated according to the standardised specific experimental framework developed by Boixel *et al*. (2019) which have been validated to be representative of *in planta* responses with respect to discrimination between cold- and warm-adapted individuals. TPCs describing *in vitro* growth rate as a function of temperature were established by fitting a quadratic function to the temperature–growth rate (or performance *P*) estimates for each strain: *P*(*T*) = *P*_*max*_ + *Curv*(*T* − *T*_*opt*_)^2^ where *Curv* is a shape parameter (Table S1 for more information on the model selection process). The key properties of TPCs were estimated through thermal traits commonly used to compare thermal sensitivities (Kingsolver, 2004; Angilletta, 2006). We have retained three parameters to describe the shape of these TPCs and quantify their characteristics: first, maximum performance (P_max_) which informs on TPC height (‘vertical shift’ modes of variation); second, thermal optimum (T_opt_) which informs on TPC position at the peak performance (‘horizontal shift’ modes of variation); third, thermal performance breadth (TPB_80_) which informs on the sensitivity of the response to temperature change around T_opt_ (‘width shift’ modes of variation). The estimates of T_min_ and T_max_ were not retained for further analysis as they fell outside the range of temperatures tested. TPC variation was assessed in two ways: (i) differences in the range and mean values of P_max_, T_opt_, and TPB_80_, assessed with parametric or non-parametric (depending on whether the assumptions of normality and homoscedasticity were verified) statistical tests for comparing variances and means; (ii) typological comparisons grouping together TPCs with similar thermal characteristics (functional thermal groups, referred to hereafter as ‘thermotypes’) based on a K-means clustering procedure applied to the covariation of P_max_, T_opt_, and TPB_80_ for all TPCs (Methods S2). The magnitude and distribution of diversity for thermal traits and thermotypes were analysed across individuals, populations, and scales, to detect differentiation in phenotypic patterns, in Chi-squared tests on the observed frequency distribution of thermotypes.

### Neutral genetic variation and population differentiation (Fig. 1 – step 3)

The 350 individuals composing the 12 *Z. tritici* populations were genotyped for 12 neutral genetic markers on DNA extracted from 50 mg of fresh fungal material from five-day cultures, following SSR amplification and sequencing in one multiplex PCR sample, and allele size annotation (Gautier *et al*., 2014; Methods S3a). Population structure was inferred with a Bayesian clustering approach under an admixture and correlated allele frequencies model implemented in STRUCTURE (Pritchard *et al*., 2000). The degree and significance of genetic variability within a population (genetic diversity and allele richness) and differentiation between populations (pairwise estimates of Weir and Cockerham’s F-statistic – F_ST_ – and hierarchical analyses of molecular variance – AMOVA) were evaluated with random allelic permutation procedures in GENETIX (Belkhir, 2004) and Arlequin (Excoffier & Lischer, 2010) software (Methods S3b-d).

### Dimensionality of mesoclimatic environmental variation (Fig. 1 – step 4)

Air temperature data for the closest weather stations within a mean 30-km radius of the eight sampling sites were retrieved from archives of global historical weather and climate data, to obtain: (i) monthly-averaged values of 1961-1990 climate normals (Norwegian Meteorological Institute, 2019); (ii) daily data over the sampling year (US National Climatic Data Center NCDC-CDO, 2019). Local variation in mesoclimatic temperatures were summarised for four climatic time windows (1961-1990 climate normals and thermal conditions for the year of sampling averaged over the calendar year, the wheat growing season, the winter/spring periods) and for five metrics (thermal mean, range, maximum, minimum and variance), giving a total of 20 thermal variables at each site. We then established a thermal niche (*i.e.* temperature conditions of a given sampling site) classification, by assessing the importance of each of these variables for discriminating between the three contrasting Köppen-Geiger climatic zones prospected, with a nonlinear and nonparametric random forest algorithm (RF; Breiman, 2001) in the ‘*randomForest*’ package of R (Liaw & Wiener, 2002). The importance of variables was compared on the basis of two metrics assessing the inaccuracy of RF zone classification if the variable concerned is not accounted for (RF mean decrease in prediction accuracy and node impurity, *i.e.* Gini coefficient).

### Testing for signatures of local adaptation (Fig. 1 – step 5)

Two steps were taken to detect genetic and phenotypic signatures of local adaptation underlying the observed differentiation between populations. First, the degree of genetic differentiation for the set of neutral markers (F_ST_ index; Weir & Cockerham, 1984) was compared with that for phenotypic traits (P_ST_ index; Leinonen *et al*., 2006). This made it possible to infer departures from neutral expectations (Merilä & Crnokrak, 2001), to determine whether thermal traits were under selection rather than subject to genetic drift (Brommer, 2011). F_ST_-P_ST_ comparisons were conducted separately for seasonal (on T_opt_) and geographic populations (on T_opt_ and TPB_80_), on the basis of sensitivity analyses assessing the robustness of the conclusions to variations in the approximation of Q_ST_ by P_ST_ (Methods S3e). Second, correlations between local climate conditions and *Z. tritici* thermal sensitivity were evaluated, to detect signatures of adaptation. Pearson correlation coefficients and their statistical significance were established for all possible combinations of thermal traits or thermotypic compositions and for the 20 spatiotemporal thermal variables defining the thermal niche of a climatic site.

## Results

### Marked interindividual variation in thermal traits at all scales

We observed a very high level of interindividual variation for the three thermal traits chosen to describe TPCs, within a range of 0.17 to 0.46 h^-1^ for P_max_ (*in vitro* growth rate), 9.6 to 25.1°C for T_opt_, and 2.8 to 30.9°C for TPB_80_, across all 350 strains. Individual thermal phenotypes are summarised in Fig. 2 and available in Dataset S1. The average metapopulation-level responses in the seasonal and geographic data sets were remarkably similar in terms of their quadratic parameters (Welch’s two-sample *t*-test, *P* > 0.05): P(T)_seasonal_ = 0.30 - 0.00077 × (T - 18.3)² *vs*. P(T)_geographic_ = 0.30 – 0.00088 × (T - 18.2)². Interindividual variation around this average TPC was greater for the seasonal than for the geographic scale, as demonstrated by the standard shift in TPC position along the *x*- and *y*-axes (Fig. 2a) and the distinctly larger density distributions of the three thermal traits at the seasonal scale (Fig. 2b-d; Levene’s test for homogeneity of variance: *P* = 0.01 for P_max_; *P* < 0.01 for T_opt_; *P* = 0.02 for TPB_80_). Interindividual variation within populations was similar at both the geographic and the seasonal scales, with equivalent variances for P_max_ (*x̅* ± 0.06 h^-1^ [SD] on average), T_opt_ (*x̅* ± 2.59 °C [SD] on average) and TPB_80_ (*x̅* ± 5.72 °C [SD] on average) within the 12 populations (Levene’s test for homogeneity of variance: *P* = 0.07; 0.51; 0.13, respectively). The populations may therefore be considered similar in terms of their individual variances for thermal traits. By contrast, they were not similar in terms of the corresponding population means, as significant differences were detected for T_opt_ and TPB_80_ (*P* < 0.05) but not for P_max_ (*P*_geographic_ = 0.09; *P*_seasonal_ = 0.75; contrary to what would be expected under the ‘warmer is better’ hypothesis in thermal biology but which is not surprising given the fact that this is a fungus with limited growth at high temperatures; Bennett, 1987).

**Fig. 2.**
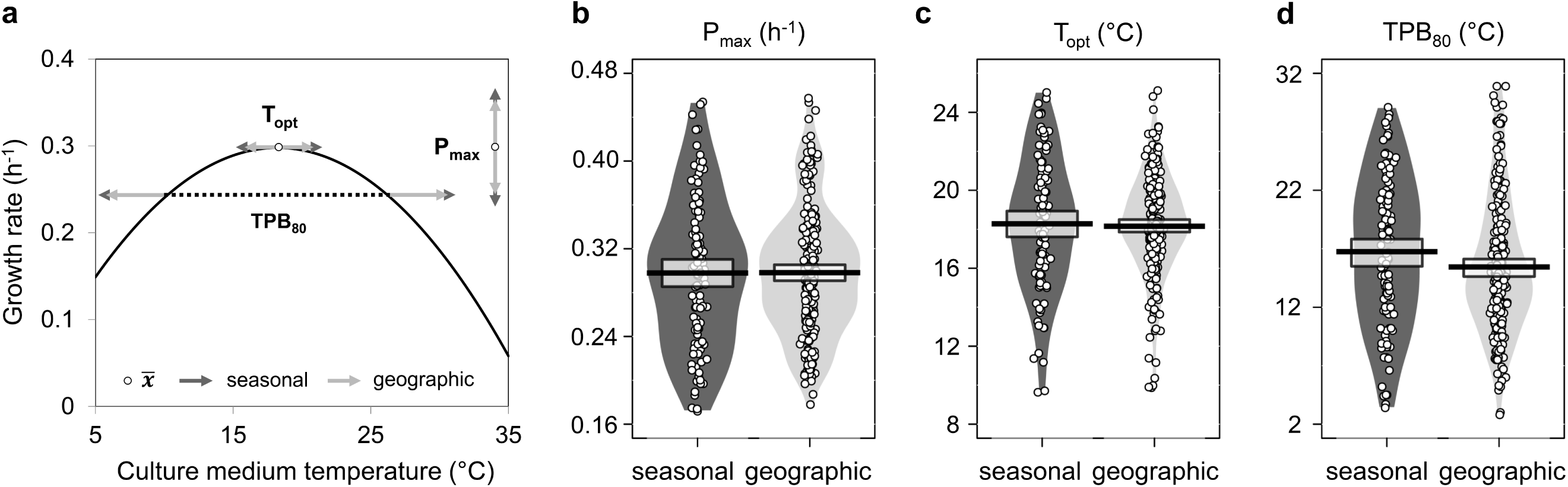
Comparisons of individual variation in *Zymoseptoria tritici* thermal performance curves (TPCs) established for *in vitro* growth rate for the seasonal and geographic scales. (a) The proportion of individual variation around the average TPC for all strains (*n* = 350) is displayed for three key thermal parameters: maximum performance (P_max_), thermal optimum (T_opt_), thermal performance breadth (TPB_80_: temperature range over which performance exceeds 80% of P_max_). The plot displays the population-level response (black solid line), the mean value over the 350 individuals for each parameter (open circles and dashed horizontal line) and the spread of the parameter (movement and shift in TPC position along the *x*- and *y*-axes) within the seasonal (*n* = 110) and geographic (*n* = 240) data sets (colour-coded arrows indicating the standard deviation around the mean). The individual variation in TPCs across strains is further broken down into the distribution of (b) P_max_, (c) T_opt_ and (d) TPB_80_, visualised as their raw individual values (open circles), means (black thick lines), distributions (smoothed density curves) and 95% Bayesian highest density intervals (central rectangular boxes enclosing the means).

### A functional reading grid for diversity in individual thermal responses

TPCs were classified into thermotypes with similar thermal responses (Hopkins’ statistic of 0.71, indicating clustered data and justifying the establishment of such a typology; Methods S2a). The diversity of TPCs encountered in the data set was optimally partitioned into 13 thermotypes (Th1 to Th13; Fig. S4), for which relative degrees of temperature specialisation were described in terms of the T_opt_ (cold- *vs.* warm-adapted), TPB_80_ (specialist *vs.* generalist), and P_max_ (low *vs.* high performer) dimensions (Fig. 3a). These thermotypes illustrated two commonly documented non-exclusive shifts in TPC along thermal gradients: a horizontal shift (low-temperature *vs.* high-temperature generalists or low-temperature *vs.* high-temperature specialists; *e.g.* Th1 *vs.* Th13 in Fig. 3b) and a generalist-specialist shift without (Th8 *vs.* Th9 in Fig. 3c) or with (Th1 *vs.* Th3 or Th11 *vs.* Th13 in Fig. 3d) trade-offs between P_max_ and TPB_80_ (*i.e.* when one cannot increase without a decrease in the other). Indeed, regression analysis revealed a significant negative correlation between P_max_ and TPB_80_ across all individuals (Pearson’s correlation coefficient: *R* = -0.44; *P* < 0.01). About 10% of individuals did not follow this pattern, with high values of both P_max_ and TPB_80_. These individuals (*i.e.* the strains of Th8), are both ‘jack-of-all-temperatures’ and ‘masters of all’, as they perform well at all temperatures (Huey & Hertz, 1984; Fig. 3e). Each cluster included strains from both geographic and seasonal populations (Fig. S4), but with an uneven distribution (difference in Jaccard distance, with a highest pairwise difference of 0.62 between WIN1 and SPR1) and an uneven relative abundance of the 13 thermotypes over the two scales. This relative abundance varied by a factor of up to two for the slightly adapted thermotypes Th5 and Th7. The various thermotypes were not equally distributed across the 12 populations either (Chi-squared test for given probabilities, *P* < 0.01). This heterogeneous distribution was particularly pronounced for high-temperature generalists (see the contributions of Th12 and Th13 to the total Chi-squared score for the comparison of distributions across seasonal and geographic populations in Fig. S5d and S6c).

**Fig. 3.**
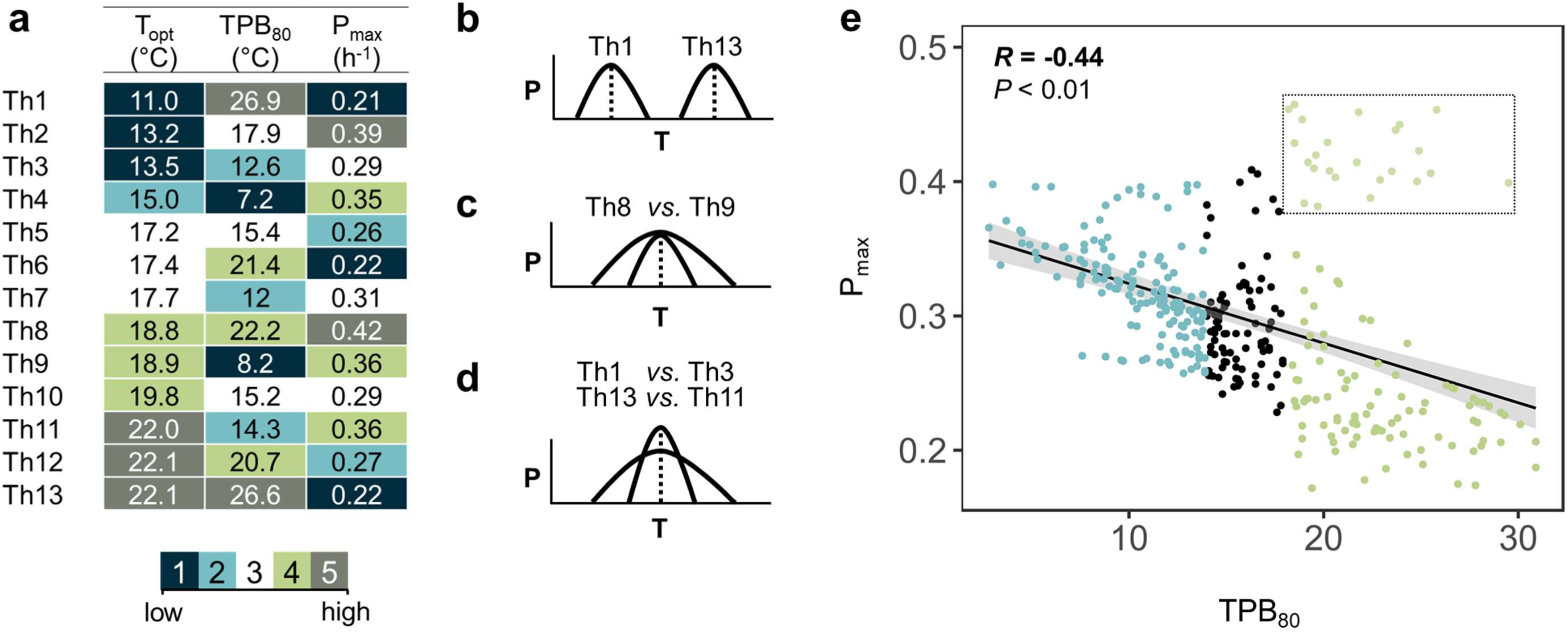
Analysis of the functional differences in thermal performance curves (TPCs) across *Zymoseptoria tritici* strains. (a) Heatmap highlighting the intrinsic features of the 13 *Z. tritici* thermotypes defined on the HCPC clustering of the 350 individual TPCs (see Fig. S4). A five-level scale was defined to summarise the overall difference in low and high values of P_max_ (low *vs.* high performance strains); T_opt_ (cold-*vs.* warm-adapted strain); TPB_80_ (specialist *vs.* generalist strain): statistically significant (1) much lower, (2) lower, (3) no deviation, (4) higher, (5) much higher value, relative to the overall mean of each parameter over the whole data set. The indicated T_opt_, TPB_80_ and P_max_ values correspond to the ‘barycentre’ of each thermotype. (b,c,d) Three common documented shifts in thermal biology studies were identified: (b) a horizontal shift with variations in the position of TPCs along the temperature-axis distinguishing ‘cold-adapted’ *vs.* ‘warm-adapted’ thermotypes; a horizontal stretch distinguishing ‘generalist’ *vs.* ‘specialist’ thermotypes (c) without or (d) with trade-offs between P_max_ and TPB_80_ (TPC axes: P: Performance; T: Temperature). (e) Scatter plot highlighting a trade-off between P_max_ and TPB_80_. P_max_ is generally negatively related to TPB_80_ except for a group of TPCs with both high TPB_80_ and P_max_ (green points surrounded by a rectangle). The regression is displayed as a solid line, with its 95% confidence interval as a shaded area, together with the Pearson’s correlation coefficient *R* and its *p*-value *P*.

Four thermotypes together accounted for almost half the entire data set (Th5, Th6, Th7 and Th10). The distinguishing features of these four thermotypes were their average behaviour with respect to T_opt_ (Th5, Th6, Th7), TPB_80_ (Th5, Th10) and P_max_ (Th7, Th10).

### Thermal phenotypic differentiation of Euro-Mediterranean populations

For population-level TPCs, significant variation was observed for thermal trait means for T_opt_ (Kruskal-Wallis, *P* < 0.01) and TPB_80_ (Kruskal-Wallis, *P* < 0.01), but not for P_max_ (Kruskal-Wallis, *P* = 0.09), for which no population differentiation was detected (Table 2). P_max_ values may have been constrained by the detection thresholds for optical density (smoothing of data for individuals with ‘extreme performance phenotypes’). A significant horizontal shift in T_opt_, by about 2°C compared to the 7 other populations, was observed for the IS population (Table 2), which consisted of individuals performing best at higher temperatures (Fig. 4a). Indeed, the proportion of high-temperature generalists (Th12 and Th13) was higher in the IS population (1:3 vs. 1:15 on average for the other geographic populations), accounting for 20.7 % of the imbalance in the distribution of thermotypes between populations (see contributions to the total Chi-squared score in Fig. S5d). The thermotypes best adapted to colder conditions (CA^+^, Th1-Th2-Th3) were particularly abundant in the Dfb populations (RU-KZ-LV), as shown by their long-tailed distributions skewed towards lower temperatures (with 6 highlighted individuals in Fig. S5a presenting a T_opt_ of about 10.4 ± 0.7°C, *i.e.* about 7°C below the mean value). The IS population was characterised by a higher TPB_80_ for its average population response than the other populations, particularly DK (19.5 *vs.* 12.7°C; Table 2). These two populations had opposite patterns in terms of their respective proportions of thermal specialists and generalists (Fig. 4b and Fig. S5b). More broadly, the individuals with the greatest thermal breadth (G^+^, Th1-Th13) were less abundant in Cfb populations (DK-FR-IR), which were characterised by a higher proportion of more highly specialist individuals (S^+^, Th4 and Th9) than the average (accounting for 10% of the total Chi-squared score; Fig. S5d).

**Table 2.** Differentiation in the averaged thermal performance curves (TPCs) of the 12 Zymoseptoria tritici populations. Each population is characterised by the population mean values for maximum performance (P_max_), thermal optimum (T_opt_) and thermal performance breadth (TPB_80_) of individual TPCs. Significant differences in the parameters of TPCs between populations were assessed separately for geographic and seasonal populations, through mean comparisons. The Latin (geographic analysis) and Greek (seasonal analysis) letters in brackets indicate significant differences in post-hoc tests.

| Thermal trait |  | Geographic scale |  |  |  |  |  |  |  | Seasonal scale |  |  |  |
| --- | --- | --- | --- | --- | --- | --- | --- | --- | --- | --- | --- | --- | --- |
|  |  | RU | KZ | LV | DK | FR | IR | TN | IS | WIN1 | SPR1 | WIN2 | SPR2 |
| $P_{\max}$ | (h<sup>-1</sup>) | 0.31 | 0.29 | 0.29 | 0.32 | 0.31 | 0.28 | 0.29 | 0.29 | 0.31 | 0.3 | 0.3 | 0.29 |
| $T_{\text{opt}}$ | (°C) | 17.1<br>(a) | 17.8<br>(a) | 17.8<br>(a) | 17.7<br>(a) | 18.1<br>(a) | 18.2<br>(a) | 18.5<br>(a) | 20.0<br>(b) | 15.9<br>(α) | 20.9<br>(β) | 17.0<br>(α) | 19.3<br>(β) |
| $TPB_{80}$ | (°C) | 15.4<br>(b) | 14.9<br>(bc) | 16.4<br>(b) | 12.7<br>(c) | 14.1<br>(bc) | 14.9<br>(bc) | 15.7<br>(b) | 19.5<br>(a) | 14.8 | 18.9 | 17 | 16.4 |

**Fig. 4.**
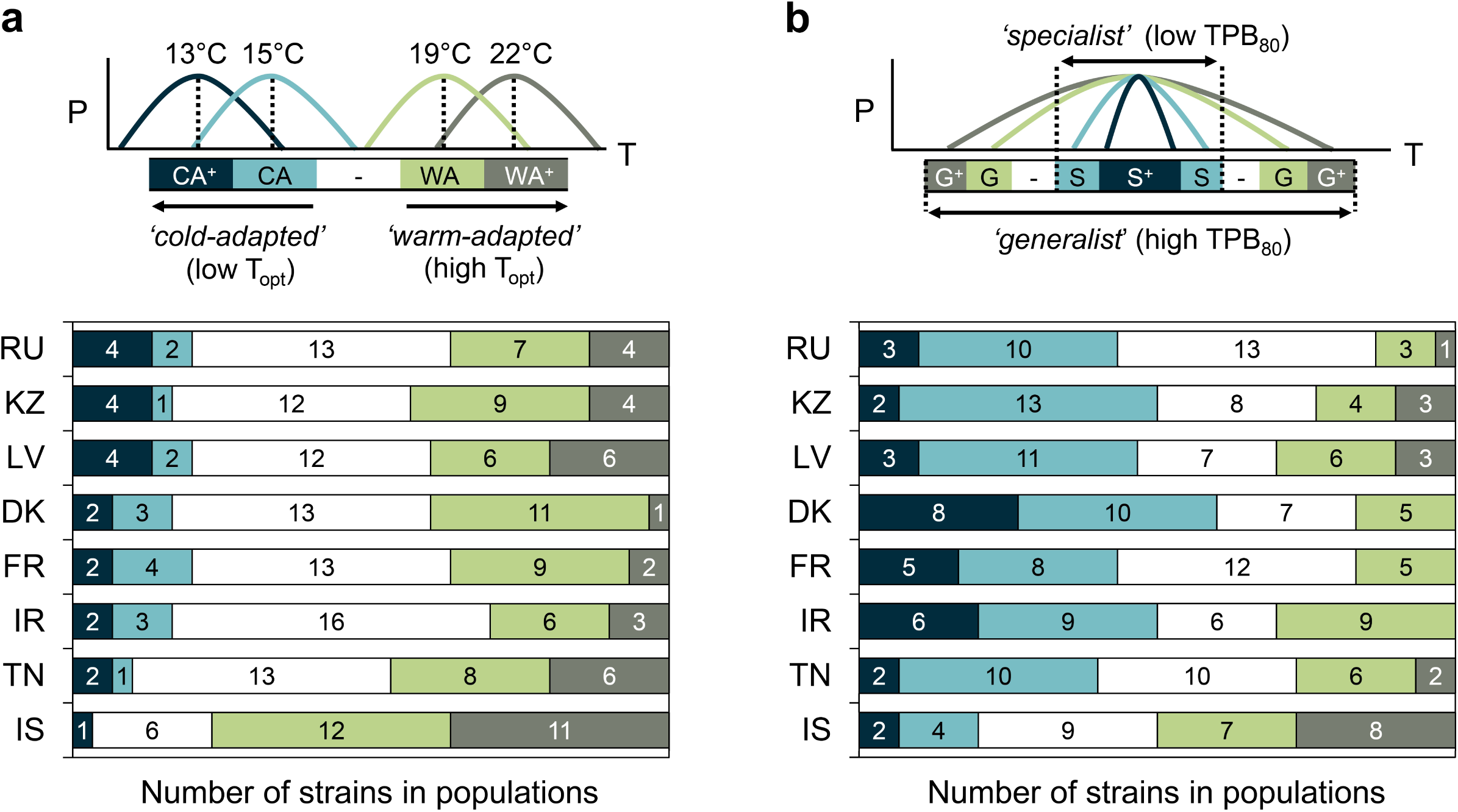
Thermal differentiation in the functional composition of the 8 geographic *Zymoseptoria tritici* populations. The functional composition of these populations is displayed according to two complementary reading grids relating to: (a) optimal temperature, with the relative proportions (*x*-axis) and corresponding number of individuals (bar values) of highly cold-adapted (CA^+^), cold-adapted (CA), intermediate (- in white), warm-adapted (WA), and highly warm-adapted (WA^+^) thermotypes within each population; (b) thermal breadth with the relative proportions (*x* -axis) and corresponding number of individuals (bar values) of high (S) and very high (S^+^) specialist (mean TPB_80_ of 10.8°C) *vs.* high (G) and very high (G^+^) generalist (mean TPB_80_ of 23.6°C) thermotypes Populations were sampled in RU (Russia), KZ (Kazakhstan), LV (Latvia), DK (Denmark), FR (France), IR (Ireland), TN (Tunisia), IS (Israel).

### Seasonal adaptive shifts within local populations

Spring subpopulations (SPR1 and SPR2) had a higher thermal optimum than winter subpopulations (ANOVA, *P* < 0.01), with a horizontal shift of T_opt_ towards higher temperature of the order of 5°C (SPR1) and 2.3°C (SPR2) on average (Table 2 and Fig. 5a). In terms of thermotype composition, these two pairs of subpopulations differed principally in their relative proportions in strains highly adapted to warm conditions (WA^+^). WA^+^ strains were significantly more abundant in SPR populations (Fig. 5b) than in WIN populations, accounting for 33.4% of the total Chi-squared score for difference in thermotype distributions between WIN and SPR populations. Conversely, WIN populations had a higher proportion of individuals highly adapted to cold conditions (CA^+^; Fig. S6).

**Fig. 5.**
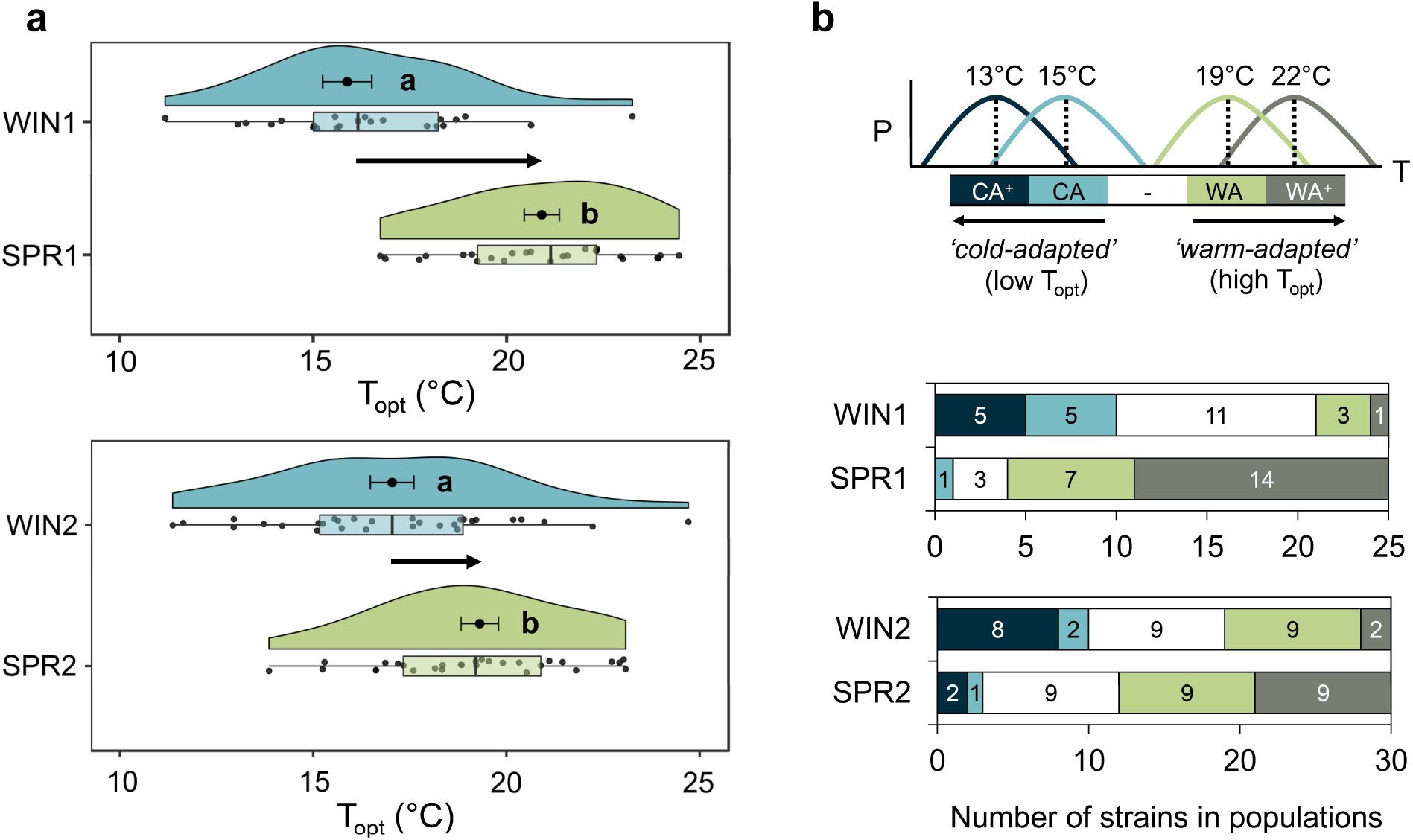
Individual differentiation in the thermal optimum of *Zymoseptoria tritici* strains between French winter and spring subpopulations. (a) The population-level thermal optima (means ± SEM) are presented together with the distribution of individual T_opt_ within populations (associated raw data points, boxplots and split-half violins). A significant shift in T_opt_ distribution along the temperature-axis was detected between winter (WIN1 and WIN2) and spring (SPR1 and SPR2) subpopulations sampled from two local neighbouring fields (annotated 1 and 2). The letters indicate the output of paired Student’s *t-*tests with *P* < 0.05. (b) Functional thermotype composition within winter and spring subpopulations is displayed as relative proportions (*x*-axis) and corresponding numbers of individuals (bar values) for highly cold-adapted (CA^+^), cold-adapted (CA), intermediate (-in white), warm-adapted (WA), and highly warm-adapted (WA^+^) thermotypes.

### Signatures of local adaptation to mean annual temperature conditions

Neutral molecular markers revealed that all strains were genetically different. We observed no difference in the genetic structure of the 12 populations, with similar allele frequencies at each locus (Fig. S7 and Table S2), suggesting a constant mixing of populations through substantial continental gene flow as underlined in previous studies for *Z. tritici* (Boeger *et al*., 1993). Strong evidence of local adaptation was detected, with the occurrence of strong phenotypic divergence (Fig. 6) and a robust P_ST_ - F_ST_ difference for the T_opt_ of both geographic and seasonal populations and for the TPB_80_ of geographic populations (Fig. S8). An analysis of possible correlations between these thermal traits and the temperature conditions of the eight sampling sites (monthly averaged values of 1961-1990 climate normals) indicated that the mean thermal optimum of geographic populations increased with mean annual temperature (Fig. 7a). The level of cold adaptation of these populations (measured as the ratio of highly cold-adapted to highly warm-adapted strains) was negatively and significantly correlated with the same environmental variable (Fig. 7b). The mean annual temperature over the 1961-1990 period to which populations seemed to be adapted was one of the three thermal variables most strongly differentiating between the three Köppen-Geiger climatic zones considered in this study (highest random forest mean decrease in accuracy; Fig. S9), together with mean annual temperature over the year of sampling and seasonal contrasts. This difference between mean spring and mean winter temperatures gave the highest mean decrease in Gini index (0.27 *vs*. 0.25 for mean annual temperature over the 1961-1990 period). This finding highlights the potential importance of seasonal conditions in structuring the thermal responses of these geographic populations.

**Fig. 6.**
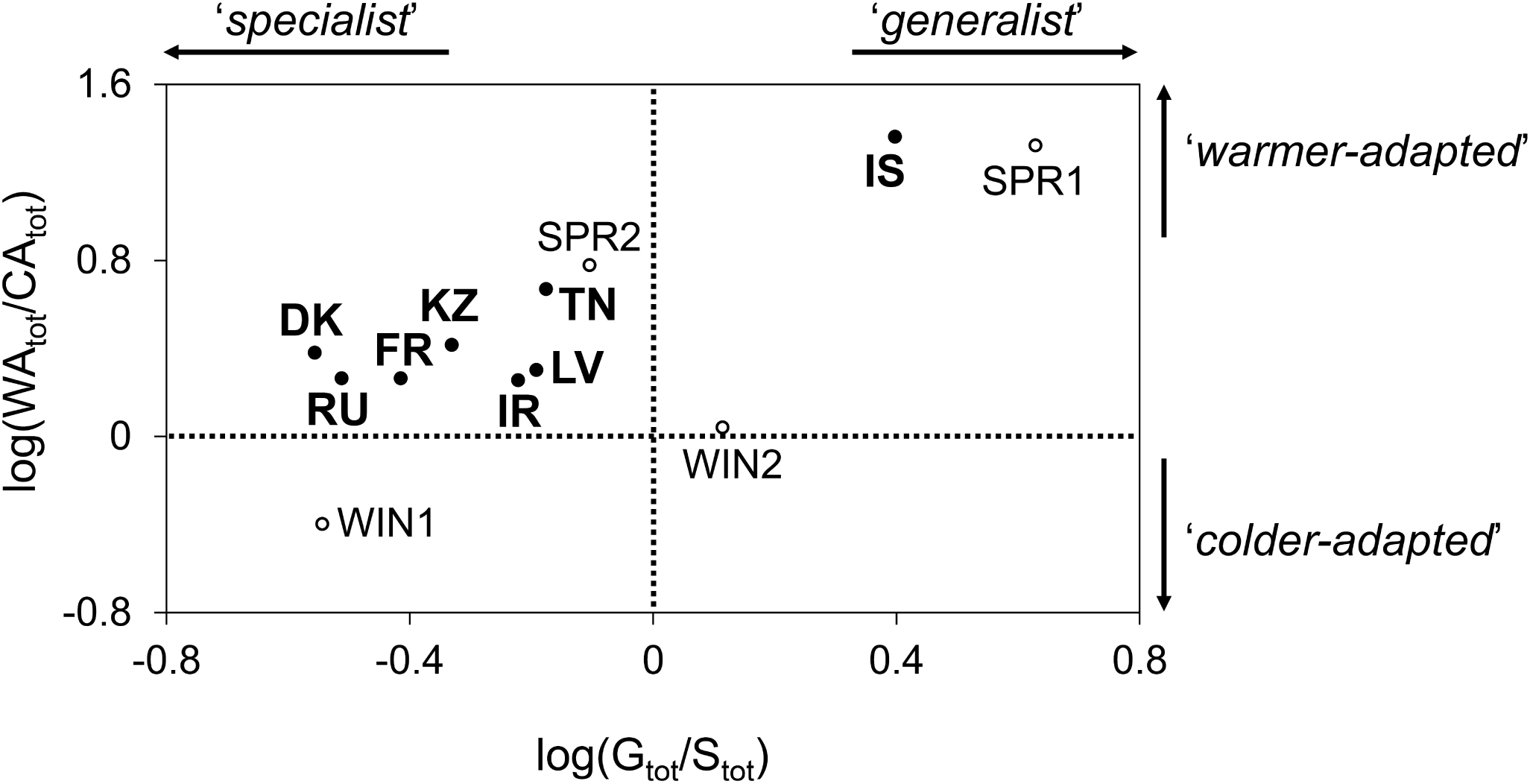
Functional diversity in thermal responses between the 12 *Zymoseptoria tritici* populations. Geographic (in bold) and seasonal (in standard text) populations are situated along: (i) a scale of increasing degree of adaptation to warm conditions (*y*-axis) discriminating colder- and warmer-adapted populations (logarithm of the ratio of the total of warm-adapted individuals – WA and WA^+^ – to the total of cold-adapted individuals – CA and CA^+^); (ii) a scale of thermal breadth continuum (*x*-axis) discriminating more specialist and more generalist populations (logarithm of the ratio of the total number of generalist individuals – G and G^+^ - to the total number of specialist individuals – S and S^+^).

**Fig. 7.**
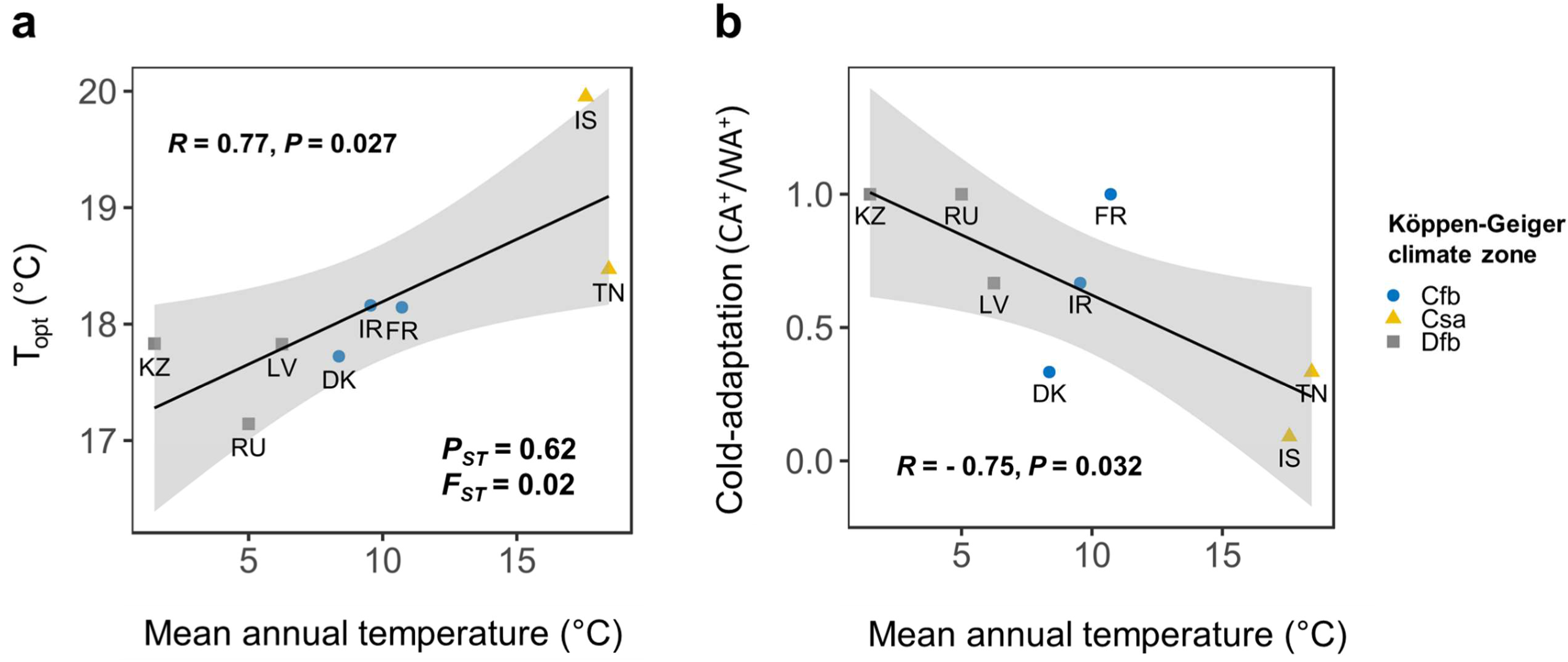
Signatures of *Zymoseptoria tritici* adaptation to the mean annual temperature of the local environment in the 8 geographic populations. (a) Relationship between population thermal optimum (T_opt_) and the mean annual temperature of the sampling sites (monthly averaged values of 1961-1990 climate normals, themselves positively correlated with the monthly averaged values over the sampling year; Pearson’s correlation coefficient: *R* = 0.98, *P* < 0.01). Population differentiation in T_opt_ relative to neutral genetic differentiation is indicated by P_ST_ and F_ST_ values. (b) Relationship between cold-adaptation level, defined as CA^+^ / WA^+^ (ratio of the number of highly cold-adapted to highly warm-adapted thermotypes), and the mean annual temperature at the sampling site. Linear dependence between these pairs of variables is indicated by the regression line (solid line and its 95% confidence interval, shown as a shaded area), Pearson’s correlation coefficient *R* and its associated p-value *P* (see Fig. S1 for a description of the three Köppen-Geiger climate zones, Cfb, Csa and Dfb).

## Discussion

### Striking spatiotemporal diversity and distribution of *Z. tritici* thermal responses

By characterizing the TPCs of *Z. tritici* strains collected over different spatiotemporal scales, we were able to develop a fine description of the extensive interindividual variation in thermal plasticity: maximum performance (P_max_), thermal optimum (T_opt_), and thermal performance breadth (TPB_80_). This detailed characterisation was made possible by the large range of temperatures and the high resolution of this experimental study (12 temperatures, ranging from 6.5 to 33.5 °C), the extensive sampling strategy (350 strains from 12 populations collected within the Euro-Mediterranean region) and the use of a dedicated and previously validated experimental framework based on turbidity measurements (Boixel *et al*., 2019). It is important to bear in mind that these turbidity measurements may not reflect the sole growth multiplication rate *via* yeast-like budding but more precisely quantify the total fungal biomass that could be affected by the pleomorphic nature of some strains of *Z. tritici* under some environmental stimuli (*e.g.* partial transition to pseudohyphae or induction of a few chlamydospores, a very recently highlighted form at high temperatures; Francisco *et al*., 2019). Precautions were taken to work under culture conditions limiting morphological transitions in the four-day time window of the experiments: very few hyphae were observed at 96 h when validating the method (see ESM1-3 in Boixel *et al*., 2019), and no chlamydospore has so far been described in the literature before 96 h in liquid medium (Francisco *et al*., 2019). Furthermore, effects of these potential – although undetected here – morphological transitions on the estimation of thermal parameters can be neglected as it results from a double integration based on kinetics, which mathematically limits the impact of the latest time point measurements that are the most likely to be affected by morphological transitions. As such, this framework enables to detect differences in thermal sensitivity between isolates (whatever the physiological bases that underpin these differences) and to go beyond the usual tests of ‘thermal sensitivity’ based on two temperatures, which can be misleading due to the non-linearity of reaction norms (Angiletta, 2009). The interindividual variation of thermal traits was conserved across populations (similar variance within populations) but was generally more marked over the seasonal scale (for a similar average metapopulation-level response between seasonal and geographic scales). These findings are particularly striking because the choice of geographic populations made it possible to cover three contrasting Köppen-Geiger climatic zones (Fig. S1).

### Singular geographic patterns of *Z. tritici* population adaptation to local conditions

The geographic variation of TPCs provides evidence of thermal adaptation to local conditions in *Z. tritici,* with: (i) an increase in the mean thermal optimum of a given population with the annual mean temperature of its location of origin; (ii) a particularly marked adaptation to high temperatures of the population sampled in Israel, consistent with the results obtained for another Israeli population investigated by Zhan & McDonald (2011); (iii) differences in the level of specialisation of individuals between populations with higher proportions of specialist individuals in the Cfb (climatic zone with lower annual temperature range) than in the Dfb (climatic zone with higher annual temperature range) populations, consistent with the assumption that thermal generalists are favoured in more variable environments. By contrast, over a smaller geographic scale (France), using the same experimental method, we detected: (i) high levels of local diversity but no structuring of thermal responses between spring populations sampled along a gradient of increasing mean annual temperature; (ii) a marked difference between post-winter populations sampled along a gradient of increasing annual temperature range: presence of thermal generalists in the population exposed to the largest annual temperature range (19.9°C) vs. the complete absence of such generalists in the population exposed to the smallest annual temperature range (11.9°C; Boixel *et al*., 2019). The phenotypic differentiation of thermal responses at the population level probably results from local short-term selection of the fittest strains over the course of an annual epidemic. We investigated the adaptation to the location of origin of populations with respect to mesoclimatic temperature conditions. The patterns of adaptation detected may have have been blurred by a non-optimal descriptive resolution of the thermal niche. Indeed, the microenvironment actually perceived by organisms can diverge from the surrounding macroenvironment due to complex biophysical filters across scales (here phylloclimate *vs.* mesoclimate; Chelle, 2005). In a second approach scaling the actual climate perceived by *Z. tritici* populations down to the phylloclimate would help refining the definition of a thermal niche for each population (Pincebourde & Woods, 2012; Pincebourde & Casas, 2019). Such an approach might provide deeper insight into the maintenance of high levels of diversity and some degree of maladaptation in individual thermal responses within each population.

### Seasonal dynamics of thermal responses in two local *Z. tritici* populations

Sampling over the geographic scale occurred during spring, between the two time points investigated at the seasonal scale (*i.e.* post-winter and post-spring conditions). These seasonal samplings highlighted a marked seasonal shift of TPCs towards higher temperatures and changes in the thermotype composition of two local *Z. tritici* populations. This result is consistent with previous observations of seasonal short-term selection on aggressiveness traits (Suffert *et al*., 2015). This study thus reveals a two-tier thermal adaptation, with seasonal dynamics nested within and potentially occurring in each geographic local adaptation over annual epidemics. This key finding shows that adaptive patterns are ‘eco-evolutionary snapshots’ that should be interpreted with caution, to such an extent that certain evolutionary dynamics of microbial populations can be of one type over a very short time scale and another type over longer time scales. Indeed, adaptive dynamics may differ with the time scale investigated (annual or pluriannual), particularly for annual crop pathogens with both sexual and asexual reproduction cycles, such as *Z. tritici* (Suffert *et al*., 2018). Our findings could be summarised by the counterintuive statement ‘local seasonal adaptation is stronger but more fleeting than geographic adaptation’ although we would expect that regions with lower seasonal contrasts in temperature (*e.g.* with mild winters) will exert weaker selective pressure.

The use of sequential temporal sampling would make it possible to capture shifts in thermal adaptation over and between wheat-growing seasons and to detect potential trade-offs between aggressiveness and survival over winter (*e.g.* Montarry *et al*., 2007).

### From adaptation patterns to eco-evolutionary processes

Consistent with previous studies, our findings highlight the existence of high levels of genetic diversity and an absence of its structuration across *Z. tritici* populations collected from local wheat fields (Zhan *et al*., 2001) up to the regional and continental scales (Schnieder *et al*., 2001; Linde *et al*., 2002) or over the course of an epidemic cycle (Chen *et al*., 1994; Morais *et al*., 2019). The high level of gene flow suggested by this low level of genetic differentiation between populations may partly explain the maintenance of some degree of maladaptation to local conditions (*e.g.* the detection of three CA+ individuals in the IS population). More generally, we observed almost all the T_opt_-adapted thermotypes (CA+, CA, WA, WA+) in each phenotyped population (except that CA individuals were absent from the IS population and CA+ individuals were absent from the SPR1 population), despite the clear patterns of adaptation observed for T_opt_ and the large differences in environmental temperatures. This maintenance of diversity suggests that *Z. tritici* is highly tolerant to thermal variations (high probability that environmental conditions are favourable to the development of at least some individuals in a given local population). One possible explanation for this finding is that the substantial adaptation of populations to their environments (*e.g.* only warm-adapted individuals under a warm environment) is hindered by a balance between gene flow and local selection (Ronce & Kirkpatrick, 2001). It also raises the issues of the occurrence of counter-selection during the interepidemic period that might explains how local populations shift in thermotype structure to reestablish similar structures between years through heritability and genetic reassortment during sexual reproduction which is driven by antagonistic density-dependent mechanisms (Lendenmann *et al*., 2016; Suffert *et al*., 2019). Further studies are required to determine the extent to which the detected pattern of geographic adaptation is driven by the thermal conditions of the environment. For this, the potential counteracting effects of selection, gene flow, random genetic drift, mutation and recombination on the increase or decrease in genetic variation would need to be assessed (Hanson *et al*., 2012). In particular, the combination of the high diversity of thermal responses in *Z. tritici* highlighted here, their heritability (Lendenmann *et al*., 2016) and the high level of local heterogeneity within wheat canopies (Chelle, 2005) suggest that local thermal conditions probably exert strong selection pressure on thermal sensitivity (for which TPCs are probably the best proxy as they may themselves be direct targets of selection; Scheiner, 1993; Via, 1993), even in the presence of high gene flow.

### Functional group composition: a browsing pattern for investigating population dynamics

Our study illustrates how the functional classification of TPCs into thermotypes with multivariate statistical procedures can provide a complementary means of deciphering diversity patterns in the biological responses quantified in reaction norms. In particular, it constitutes an operational tool for assessing functional similarity at the individual level (*i.e.* whether the apparent variation observed in thermal parameters is functionally significant; Fig. 8a and 8b) and at the population level (*i.e.* whether the thermotypes constituting a population are more or less well-differentiated within the whole functional space; Fig. 8c). However, caution will be required in the extension of this approach to comparisons over multiple data sets, through either: (i) the development of comparable classification systems, taking into account the variation of the classification with the populations sampled by explicitly stating which ranges of trait values are hidden behind a given group description (*e.g.* affixing levels of adaptation: very low, low, high, very high); or (ii) the validation of group delineations between experiments, by combining *a priori* and *a posteriori* methodologies. This description of populations in terms of functional groups makes it possible to move from a description of phenotypic patterns and shifts in population composition to an inference process. This process may, for example, be based on comparisons of the competitive advantage of thermotypes under given thermal scenarios: *e.g.* ‘do more variable environments favour thermal generalists?’ or ‘is there a shift in the optimal range of thermal responses with mean temperature conditions?’ (Fig. 8d). This classification into thermotypes enabled here to go beyond a purely descriptive framework and future investigations will need to be undertaken to tackle the physiological bases of these differentiations in thermal responses. The thought-provoking results of Francisco *et al*. (2019) could be used to test whether several strains belonging to those thermotypes also correspond to specific or main morphotypes that would increase their tolerance under some environmental conditions (*e.g.* if warm-adapted individuals exhibit higher proportions of stress tolerant growth forms such as chlamydospores under warmer temperatures). All in all, this functional approach lays the foundations for future studies of the potential of pathogen populations to adapt to changes in their environment, from seasonal changes in the short term, to global warming in the long term. In particular, it will prove useful in gaining a fuller understanding of how new aggressive fungal strains may emerge and expand into previously unfavourable environments (Milus *et al*., 2009; Mboup *et al*., 2012; Stefansson *et al*., 2013). This is a crucial area of investigation that is all too often overlooked in models for predicting plant disease epidemics in conditions of climate change (West *et al*., 2012).

**Fig. 8.**
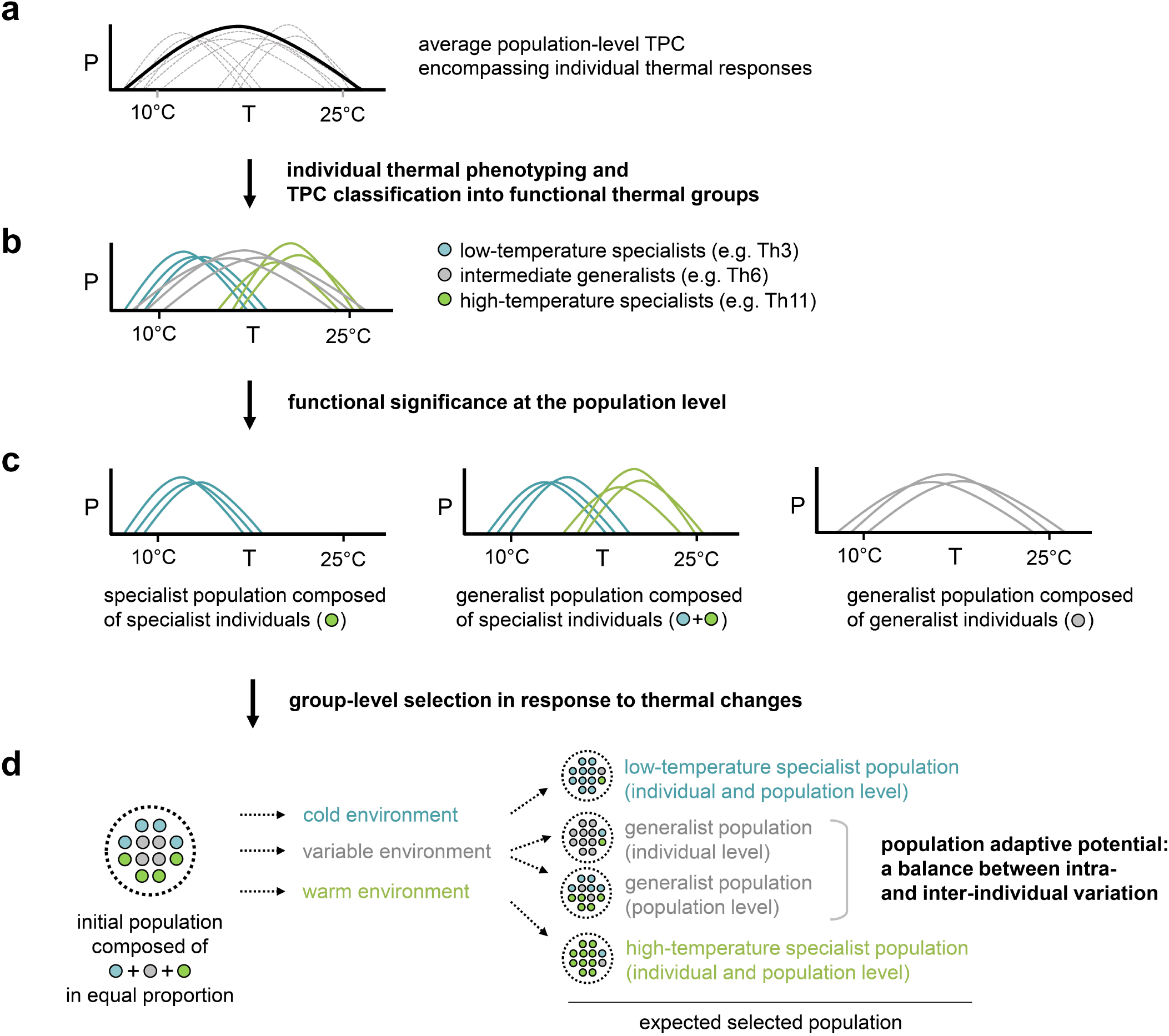
Summary of the way categorisation into thermotypes (functional thermal groups) sheds light on the translation of population diversity patterns into selection dynamics in response to climate conditions. (a) Average population-level TPC (solid line) concealing a set of varied individual TPCs (dashed lines); (b) Breakdown of the variation in individual TPCs based on their classification into thermotypes and screening for a functional significance of variation at the individual level (given example of three thermotypes within which individuals are considered functionally redundant: low-temperature specialists, intermediate generalists and high-temperature specialists). (c) Categorisation tackling functional redundancy at the population level (i.e. whether the thermotypes composing a population are more or less well-differentiated within the whole functional space). The three populations presented here demonstrate the relevance of considering functional redundancy *vs.* vacant functional space when assessing emergent properties of populations such as a generalist nature at population level (e.g. a generalist population can be composed of specialist individuals with narrow individual TPB_80_ distributed over the functional space, resulting in broad TPB_80_ population coverage). (d) The translation of populations into functional groups makes it possible to investigate group-level selection, testing for general assumptions of adaptation to given environments (e.g. competitive advantage of low-temperature specialists in cold environments, generalists in variable environments, high-temperature specialists in warm environments) providing insight into the potential of populations to adapt to changes in their environment (a subtle balance between diversity levels for intra- and inter-individual variation in thermal responses).

## Concluding remarks

We detected a high level of functional divergence in the plasticity and variation of individual thermal responses over geographic and seasonal scales, highlighting the occurrence of a two-tier dynamics in thermal adaptation. These findings raise intriguing questions regarding the mode of selection operating on these functional groups of individuals with similar competitive advantages in given thermal conditions. Deciphering the mechanisms underlying this maintenance of diversity in population phenotypic composition will prove useful for expanding our understanding of eco-evolutionary responses and the potential of populations, species and communities to adapt to environmental change.

## Acknowledgements

We would like to thank Aigul Akhmetova (*CIMMYT*, Kazakhstan), Yerlan Dutbayev (*National Agrarian University*, Kazakhstan), Andrea Ficke (*Bioforsk Plant Health and Plant Protection*, Norway), Inga Gaile (*Integrētās Audzēšanas Skola*, Latvia), Lise Jørgensen (*Aarhus University*, Denmark), Steven Kildea (*Teagasc*, Ireland), Elena Pakholkova (*Research Institute of Phytopathology*, Russia), Antonio Prodi (*University of Bologna*, Italy), Hanan Sela (*Tel-Aviv University*, Israel), Sandrine Gélisse, Thierry Marcel and Anne-Sophie Walker (*INRA BIOGER*, France) for their crucial help in collecting wheat leaves with STB symptoms, which was required for the establishment of the Euro-Mediterranean *Z. tritici* collection upon which this study is based. We would also like to thank Sylvain Pincebourde (*IRBI-CNRS*, France) for many fruitful discussions. This research was supported by a grant from the French National Research Agency (ANR) as part of the ‘Investissements d’Avenir’ programme (SEPTOVAR project; LabEx BASC; ANR-11-LABX-0034) and by a PhD fellowship from the French Ministry of Education and Research (MESR) awarded to ALB.

## Author contributions

ALB, MC and FS designed the study and wrote the manuscript. ALB performed the experiments and analysed the data.

## Supporting information

Additional Supporting Information may be found online in the Supporting Information section at the end of the article.

## Supporting Information

### FIGURES

**Fig. S1** Selection of the sampling sites in the Euro-Mediterranean wheat-growing area

**Fig. S2** Selection of the seasonal subpopulations

**Fig. S3** Appropriate sample size for estimating diversity in TPCs within *Z. tritici* populations

**Fig. S4** Clustering of *Z. tritici* strains into 13 thermotypes

**Fig. S5** Distribution of thermotypes across the eight Euro-Mediterranean *Z. tritici* populations

**Fig. S6** Distribution of thermotypes across the four French seasonal *Z. tritici* subpopulations

**Fig. S7** Genetic diversity and population structure of the 12 *Z. tritici* populations

**Fig. S8** Sensitivity analyses for the robustness of P_ST_–F_ST_ comparisons

**Fig. S9** Characterisation of the thermal niche at each sampling site

### TABLES

**Table S1** Selection of a candidate mathematical model for establishing TPCs

**Table S2** Population-pairwise genetic distance (matrix of F_ST_-values)

### METHODS

**Methods S1** Procedure for the sampling, collection and recovery of *Z. tritici* strains

**Methods S2** Definition of *Z. tritici* ‘thermotypes’ (functional thermal groups)

**Methods S3** Procedure for acquiring and analysing multilocus genotypic data

---

**Fig. S1.**
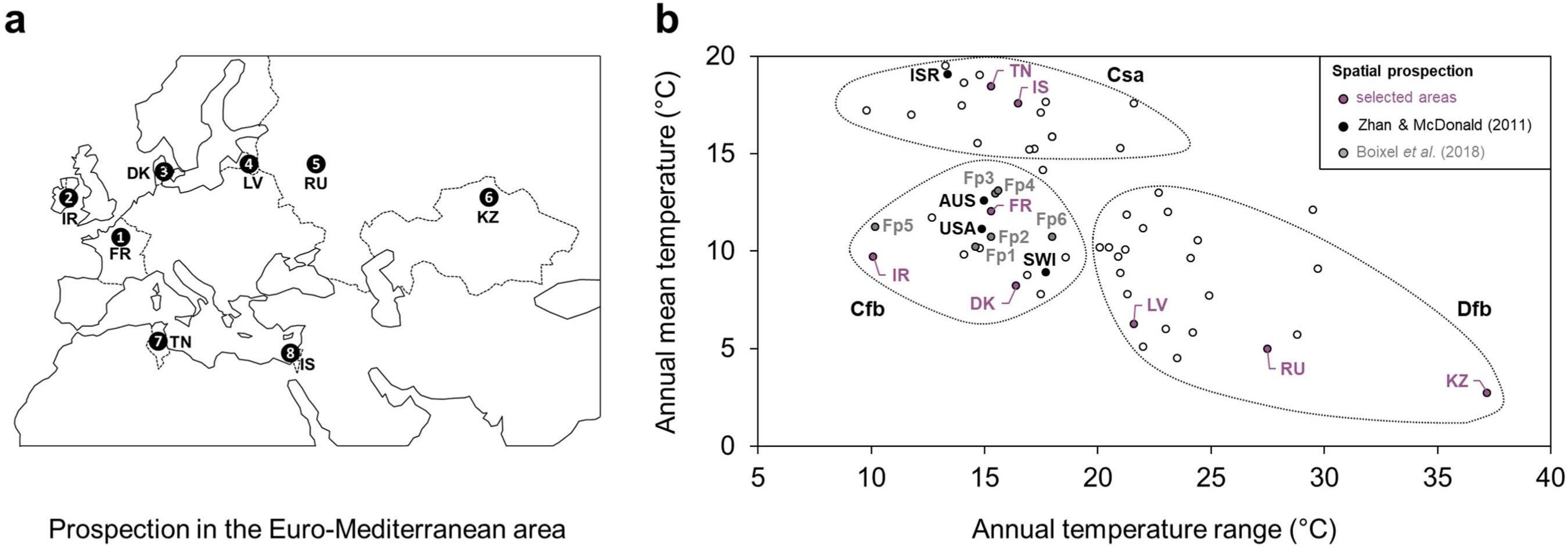
Selection of sampling sites in the Euro-Mediterranean wheat-growing area. (a) Location of the eight fields from which the Euro-Mediterranean populations of *Zymoseptoria tritici* were collected: FR (*Thiverval-Grignon*, France), IR (*Carlow*, Ireland), DK (*Flakkebjerg*, Denmark), LV (*Jelgava*, Latvia), RU (*Moscow*, Russia), KZ (*Penkovo*, Kazakhstan), TN (*Manouba*, Tunisia), IS (*Kiryat-Tivon*, Israel). Dotted lines correspond to the national borders of the countries from which the populations were collected. (b) Representative panel of climatic conditions encountered at the Euro-Mediterranean scale, for classification of the diversity of thermal responses in *Z. tritici* populations. Sampling locations (in purple) were chosen on the basis of 1961-1990 climate normals to maximize environmental heterogeneity at a level much greater than in two previous studies on thermal adaptation in *Z. tritici* (Zhan & McDonald, 2011; Boixel *et al*., 2019). The scatter plot depicts the annual mean temperature and temperature range (Norwegian Meteorological Institute, 2019) of each country location in the Euro-Mediterranean zone and in the two aforementioned studies (see open, grey and black points, respectively). The various sampling sites were classified according to their predominant Köppen-Geiger climate types (Köppen, 1936; Peel *et al*., 2007): Cfb (temperate oceanic climate), Csa (hot-summer Mediterranean climate), Dfb (warm-summer humid continental climate). AUS (*Wagga Wagga*, Australia), ISR (*Nahal Oz*, Israel), SWI (*Berga Irchel*, Switzerland) and USA (*Corvallis*, United States) correspond to the sampling locations of the five *Z. tritici* populations investigated in Zhan & McDonald (2011). Fp1 (*Villers-lès-Cagnicourt*), Fp2 (*Versailles*), Fp3 (*Bergerac*), Fp4 (*Lectoure*), Fp5 (*Ploudaniel*) and Fp6 (*Bretenière*) correspond to the sampling locations of the six French *Z. tritici* populations investigated in Boixel *et al*. (2019).

**Fig. S2.**
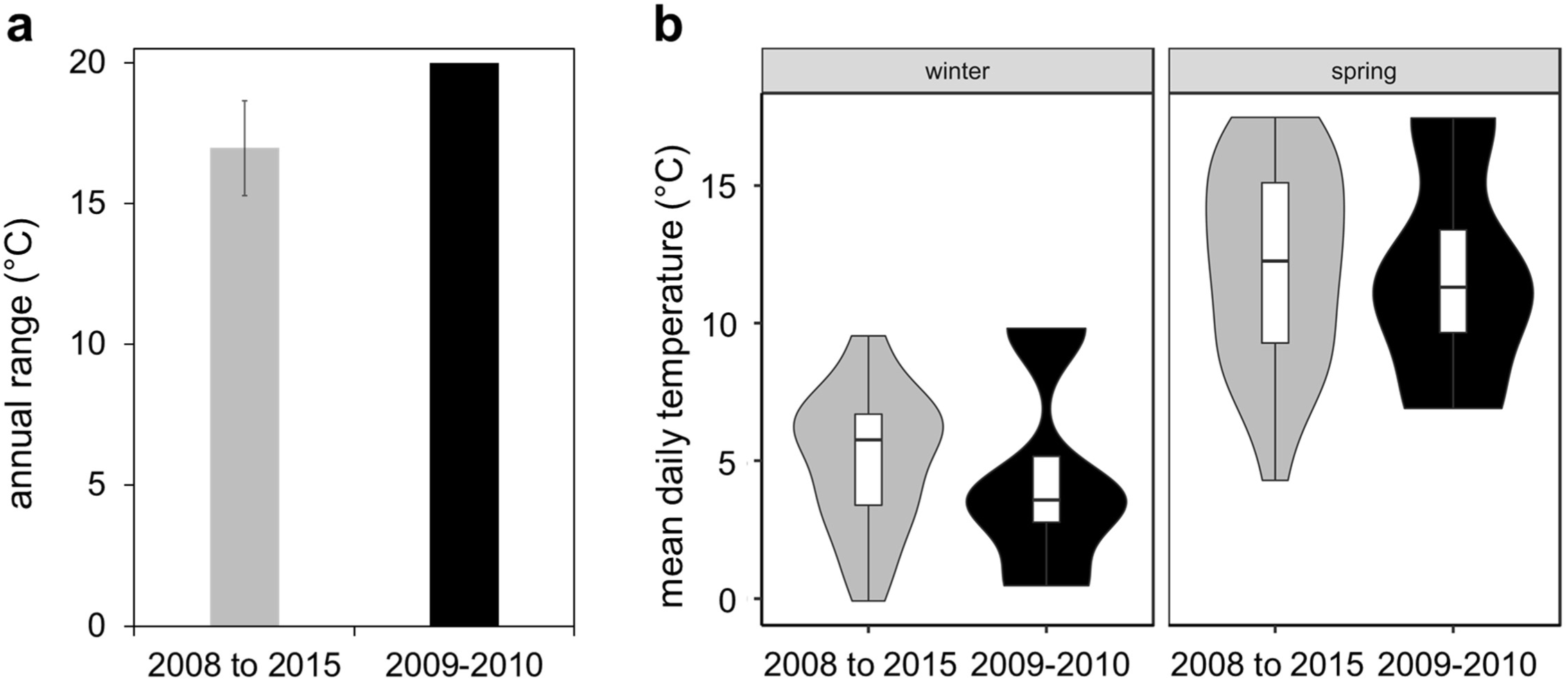
Selection of the seasonal subpopulations. Analysis of the climatic data for *Thiverval-Grignon* from 2008 to 2015 (INRA AgroClim, 2019) used to select the most contrasting wheat growing season to be investigated from those for which samples were available from the INRA BIOGER *Z. tritici* collection. Seven pairs of subpopulations were sampled from two neighbouring field plots during a long-term population survey (partially presented in Suffert *et al*., 2018) over seven wheat growing seasons (from 2008-2009 to 2014-2015): one post-winter subpopulation (sampled at the beginning of March) and one post-spring subpopulation (sampled at the end of June). We investigated seasonal effects in the 2009-2010 subpopulations, which had (a) the highest annual amplitude (20°C, vs. 16.9°C ± 1.7°C in the other seasons) and (b) the greatest contrast between ‘winter’ (November – February: tighter distribution of mean daily temperatures) and ‘spring’ (March – June: shortened low end of the distribution tail of mean daily temperatures) temperature conditions.

**Fig. S3.**
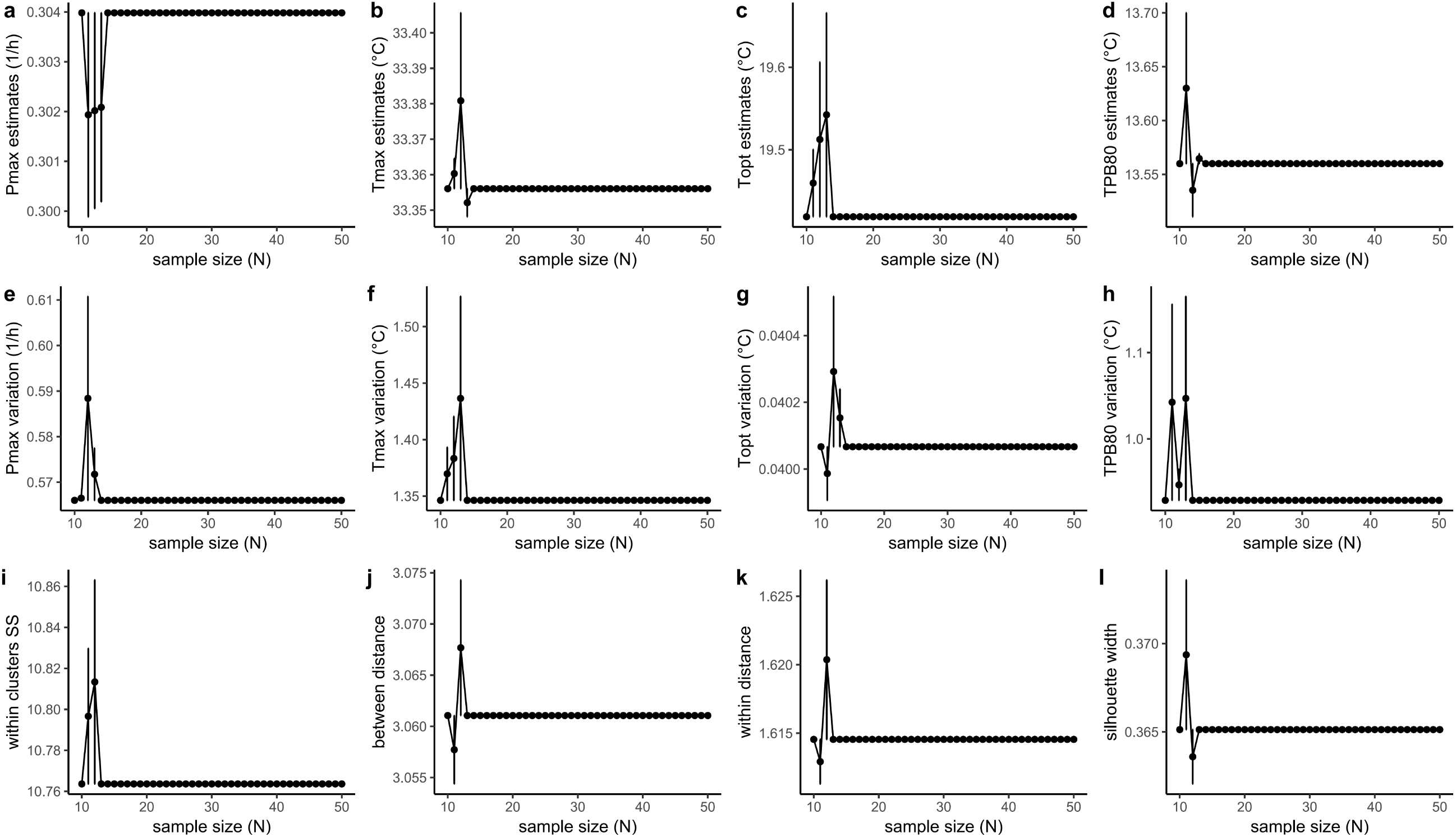
Appropriate sample size for estimating diversity in TPCs within *Z. tritici* populations. For the definition of a target sample size per population, we performed a rarefaction analysis on a set of 66 *Z. tritici* strains that had undergone preliminary phenotyping for their responses to temperature (see Boixel *et al*., 2019). Rarefaction curves of expected diversity for key thermal parameters (a: mean maximum performance P_max_; b: mean maximum temperature T_max_; c: mean thermal optimum T_opt_; d: mean 80% thermal performance breadth TPB_80_; e: P_max_ standard deviation; f: T_max_ standard deviation; g: T_opt_ standard deviation; h: TPB_80_ standard deviation) and in the typology of thermal responses based on HCPC clusters (i: within- thermotype sum of squares; j: mean distance between clusters; k: mean distance within clusters; l: mean silhouette width providing information about the compactness, separation, and connectivity of the cluster partitions) were obtained for 41 levels of sampling depth (mean ± resampling standard error; *n* = 15 subsampling repetitions per sampling depth) ranging from 10 to 50 individuals per subsample (sample size *N; x*-axis). Similar results were obtained for samples of more than 15 strains. We chose to phenotype and genotype 30 of the 50 strains that we isolated in total for each population (INRA BIOGER collection), to ensure a sufficiently high statistical power and precision and to ensure that we could estimate allele frequencies and gene diversity in a population (Dale & Fortin, 2014).

**Fig. S4.**
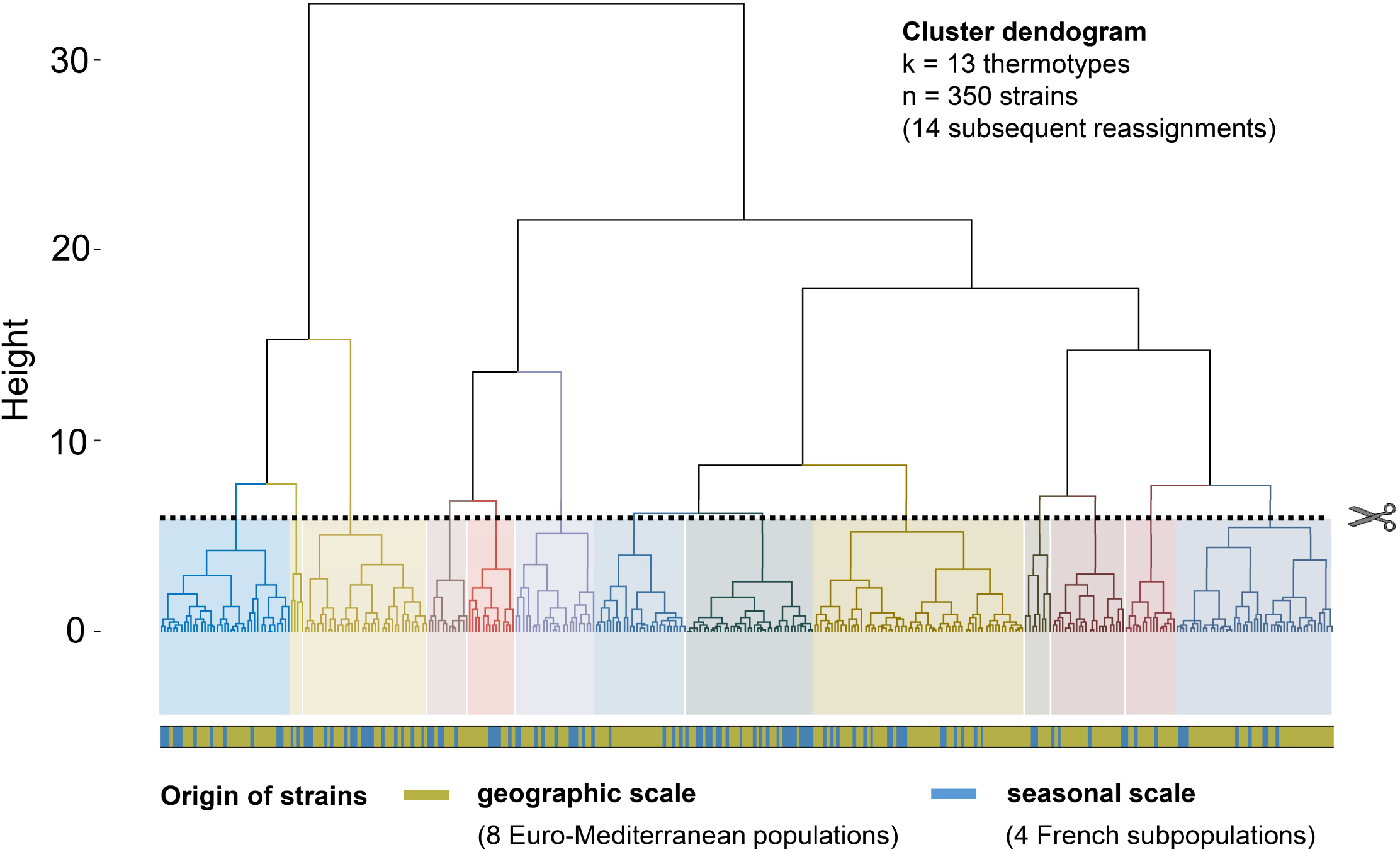
Clustering of *Z. tritici* strains into 13 thermotypes. (a) The dendogram (distance-based tree) illustrates the hierarchical organisation of the TPCs of the 350 *Z. tritici* strains (seasonal and geographic scales) in terms of functional thermal responses. The distance between strains in the hierarchical cluster tree was calculated based on Euclidean distances (the greater the difference in height, the greater the dissimilarity). We identified 13 thermotypes, colour-coded in the figure, with the procedure presented in Methods S2. Each thermotype includes strains from both data sets (seasonal: blue segments; geographic: yellow segments in the horizontal bar below the tree). These results demonstrate the relevance of establishing a unique typology to compare the TPCs of all 350 strains. The quality of the clustering was assessed by silhouette analysis (Rousseeuw, 1987), based on the mean distance between clusters. 14 individuals were placed in the wrong cluster (negative silhouette width) and were therefore reassigned to the closest neighboring cluster before assessing the distribution and features (compactness, separation, connectivity) of each thermotype.

**Fig. S5.**
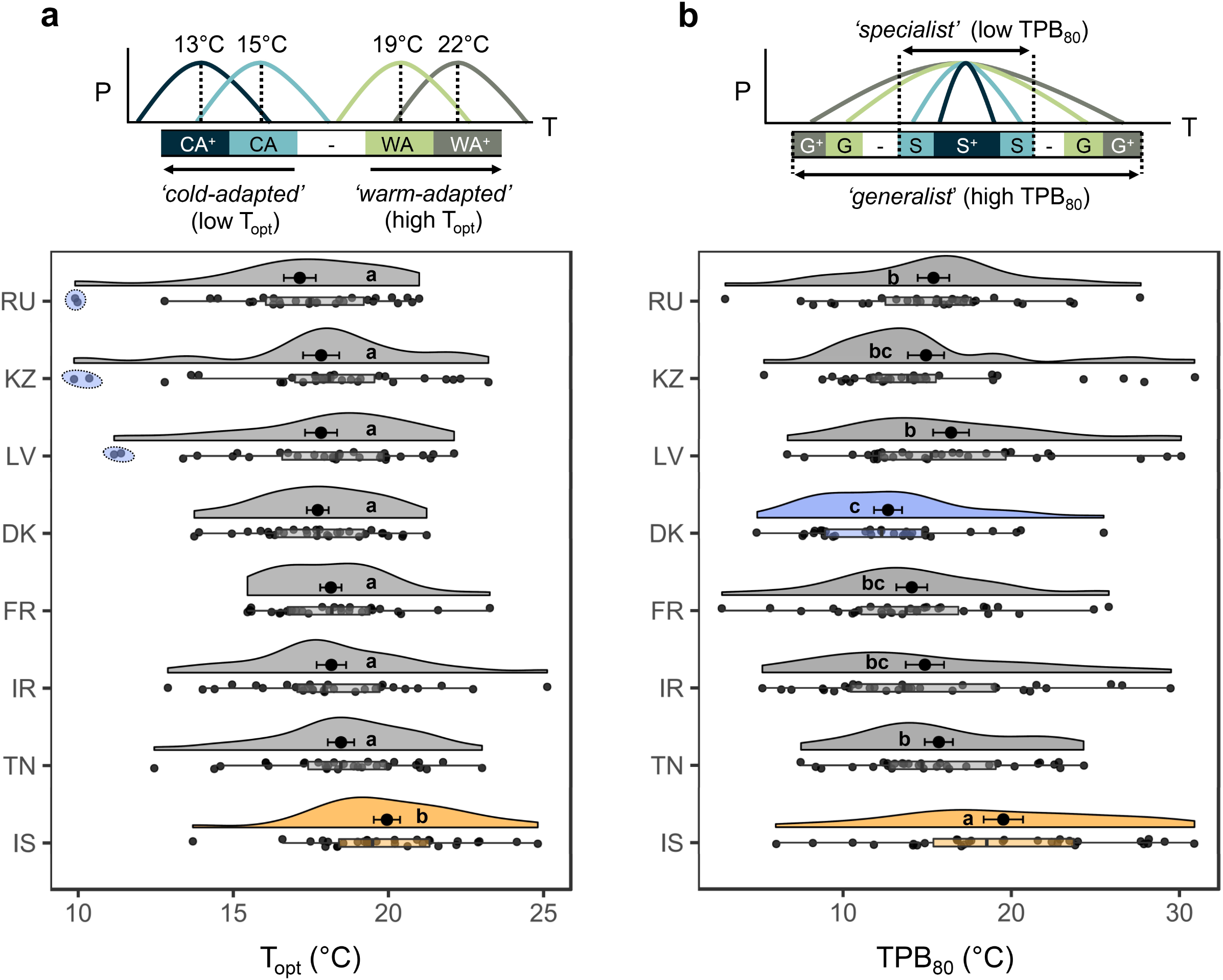
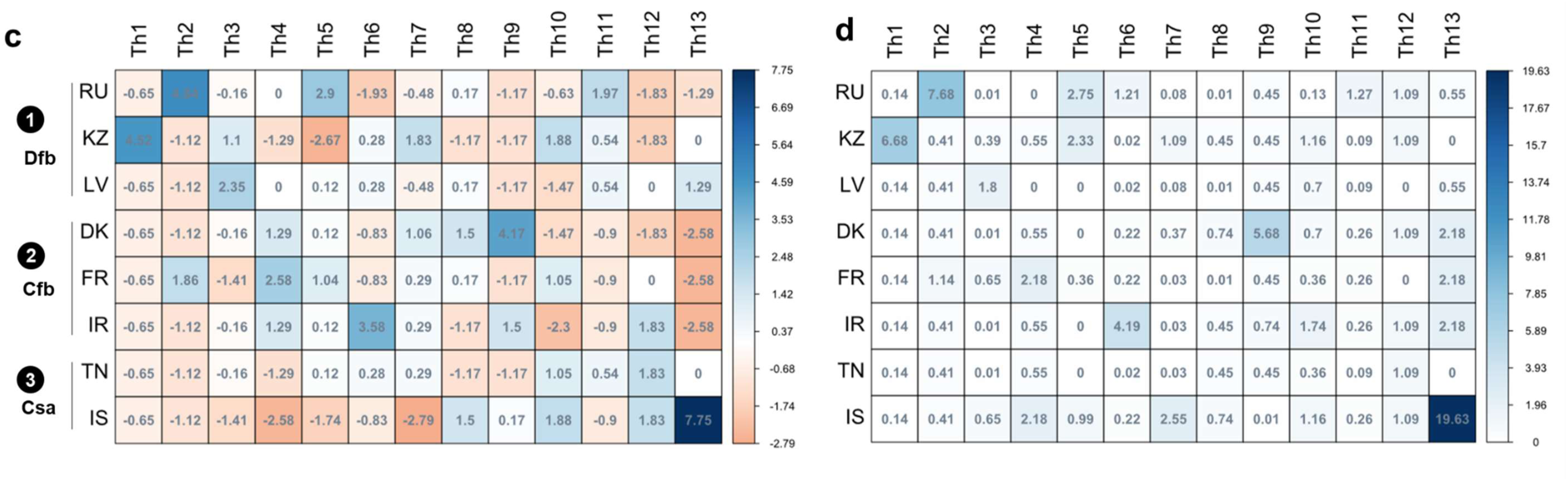
Distribution of thermotypes across the 8 Euro-Mediterranean *Z. tritici* populations. (sampled in RU: Russia; KZ: Kazakhstan; LV: Latvia; DK: Denmark; FR: France; IR: Ireland; TN: Tunisia; IS: Israel). (a) Population-level thermal optima (means ± SEM) are presented, together with the distribution of individual T_opt_ values within populations (associated raw data points, boxplots and split-half violins). A significant shift in T_opt_ distribution towards higher temperature was detected for the Israeli population (IS); the letters indicate the outputs of Kruskal-Wallis post-hoc pairwise comparisons, with *P* < 0.05. (b) Population-level thermal breadth (means ± SEM) values are presented together with the variation in that response among individuals (associated raw data points, boxplots and split-half violins). The letters indicate the outputs of Kruskal-Wallis post-hoc pairwise comparisons, with *P* < 0.05. (c) Contingency table for the Chi-squared test of independence. Each row represents a population and each column represents a thermotype. Associations between rows and columns are displayed for each cell as Pearson residuals. Positive values in blue indicate an attraction and negative values in red indicate a repulsion. This table reveals that: (1) the most strongly cold-adapted thermotypes (CA^+^, Th1-Th2-Th3) are preferentially found in Dfb populations (RU-KZ-LV); (2) individuals with the greatest thermal breadth (G^+^, Th1 and Th13) are less abundant in Cfb populations (DK-FR- IR), which are characterised by more highly specialist individuals (S^+^, Th4 and Th9) than the average; (3) warm-adapted generalist individuals (WA^+^- G/G^+^, Th12,Th13) are found in higher proportions in the IS and, to a lesser extent, TN populations. (d) The relative contribution of each cell to the total Chi-squared score (darker cells are those corresponding to the bulk of the variability in thermotype distribution) indicates that the three aforementioned aspects account for (1) 17.7 %, (2) 17.1%, and (3) 21.8% of total phenotypic differentiation in thermal response between populations.

**Fig. S6.**
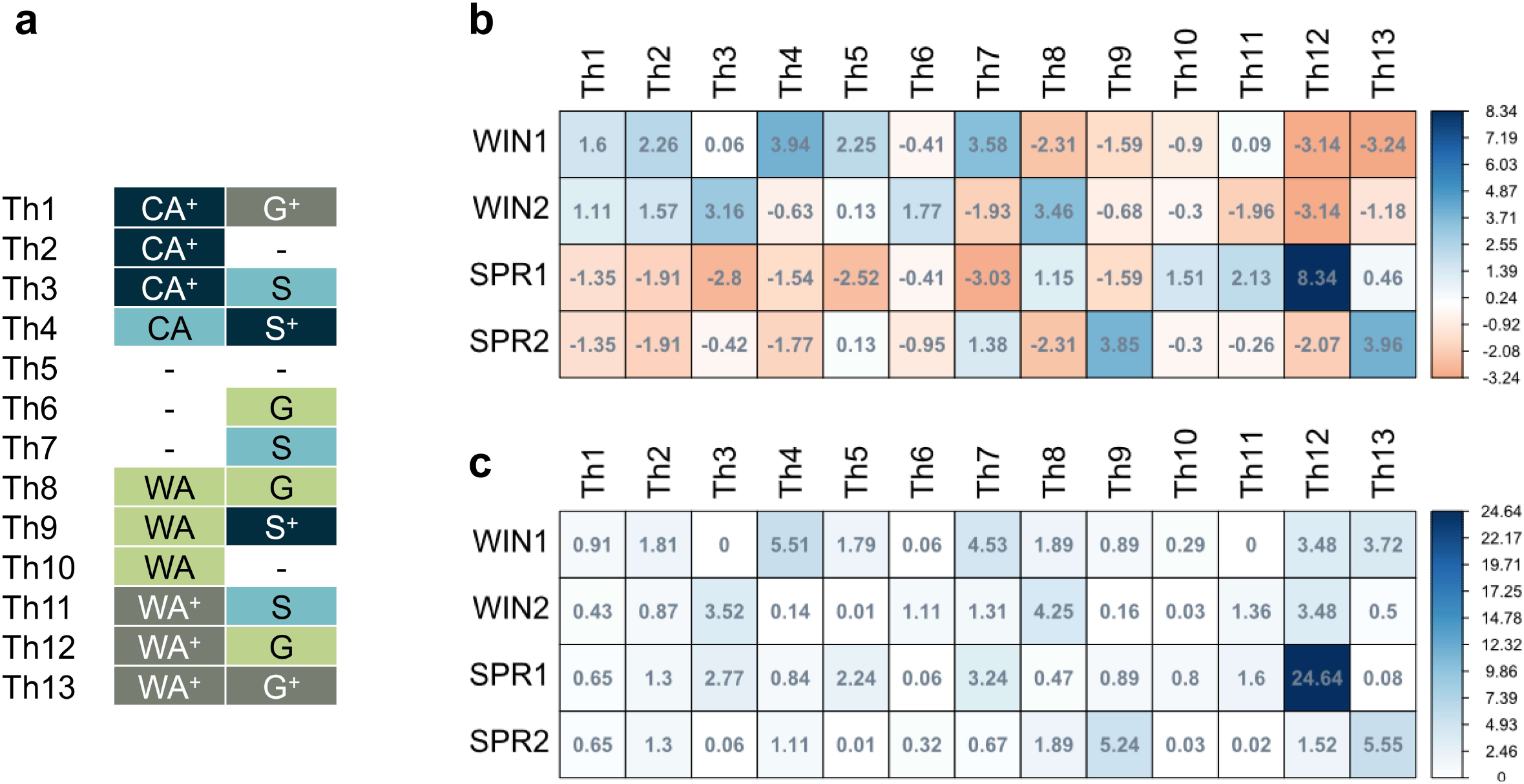
Distribution of thermotypes across the four French seasonal *Z. tritici* subpopulations. (a) Stacked bar plots of functional thermotype composition across populations: highly cold-adapted (CA^+^), cold-adapted (CA), intermediate (- in white), warm-adapted (WA), highly warm-adapted (WA^+^) thermotypes. The relative abundance within populations of each thermotype is expressed as a single bar percentage on which the thermotype ID is displayed. (b) Contingency table of the Chi-squared test of independence. Each row represents a population and each column represents a thermotype. Associations between rows and columns are displayed for each cell as Pearson residuals. Positive values in blue indicate an attraction and negative values in red indicate a repulsion. These residuals show that winter (WIN1 and WIN2) and spring (SPR1 and SPR2) subpopulations fall into discrete clusters in terms of their composition in thermotypes highly adapted to cold (CA^+^: positive association with winter populations and negative association with spring populations) and in thermotypes highly adapted to warm conditions (WA^+^: negative association with winter populations and positive association with spring populations). (c) The relative contribution of each cell to the total Chi-squared score (darker cells are those corresponding to the bulk of the variability in thermotype distribution) indicates that the compositional dissimilarity in thermotypes between winter and spring populations can be accounted for mostly by the proportions of generalist warm-adapted individuals (Th12 and Th13; 43% of the total Chi-squared score).

**Fig. S7.**
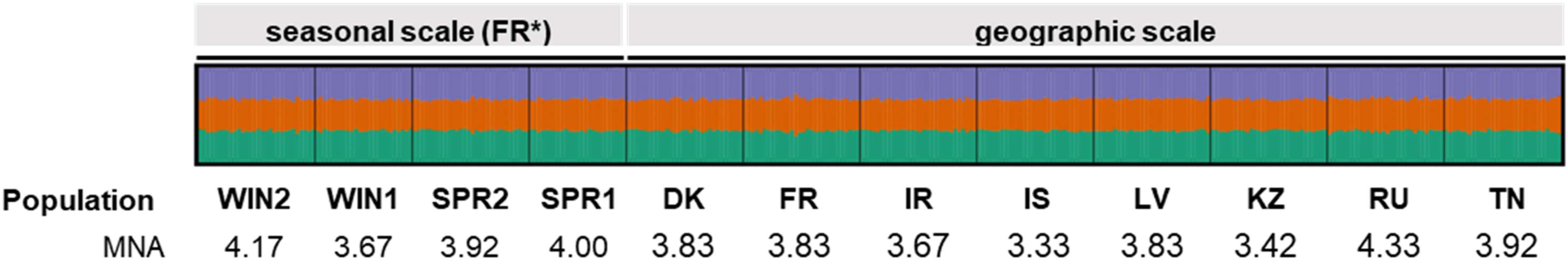
Genetic diversity and population structure of the 12 *Z. tritici* populations. The 350 *Z. tritici* strains were collected over a large spatial scale (8 geographic sites: DK: Denmark; FR: France; IR: Ireland; IS: Israel; KZ: Kazakhstan; LV: Latvia; RU: Russia; TN: Tunisia) and over a seasonal scale during another growing season in France (FR* with two pairs of post-winter (WIN1; WIN2) and post-spring (SPR1; SPR2) subpopulations). These strains were genotyped for 12 neutral microsatellite markers (SSRs) and assigned to different genetic clusters by a Bayesian cluster approach (see the detailed procedure in Methods S3). Each strain (one unique multilocus genotype) included in the analysis is displayed as a thin vertical line partitioned into colored segments representing its probabilities of assignment to genetically different clusters (cluster 1 in green, cluster 2 in orange and cluster 3 in purple). Each strain was affected to each of the three genetic structures with the same probability, indicating an absence of population structure. Genetic diversity is indicated for each population as the mean number of alleles observed over the 12 SSRs (MNA).

**Fig. S8.**
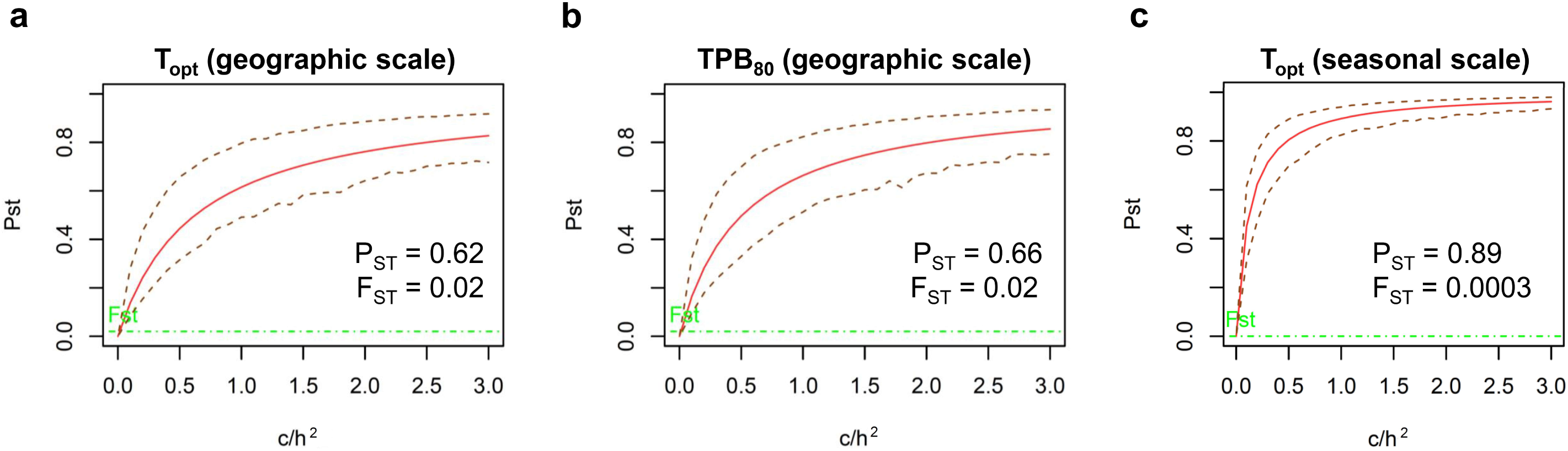
Sensitivity analysis of the robustness of P_ST_–F_ST_ comparisons for the 12 *Z. tritici* populations. The robustness of P_ST_–F_ST_ comparisons was explored by studying variations of the c/h^2^ ratio, which determines the accuracy of the approximation of Q_ST_ by P_ST_. Phenotypic divergence in thermal sensitivity (P_ST_) was investigated for: (a) the thermal optimum (T_opt_) of the Euro-Mediterranean populations (geographic scale); (b) for the thermal breadth (TPB_80_) of Euro-Mediterranean populations; (c) for the thermal optimum (T_opt_) of the French seasonal subpopulations (seasonal scale). For each plot, estimates of P_ST_ (solid line in red, the P_ST_ value displayed being calculated at the critical c/h² ratio of 1), its lower and upper 95% confidence interval limits (dashed lines in red) and the upper confidence estimate of the neutral divergence (F_ST_; dashed line in green) are plotted. For each trait, the 95% CI of P_ST_ at c/h^2^ = 1 indicates the occurrence of strong phenotypic divergence and of a robust difference in P_ST_ and F_ST_, as their confidence intervals only overlap (P_ST_ > F_ST_) at low c/h^2^ ratios (a: c/h^2^ = 0.02; b: c/h^2^ = 0.02; c: c/h^2^ < 0.005).

**Fig. S9.**
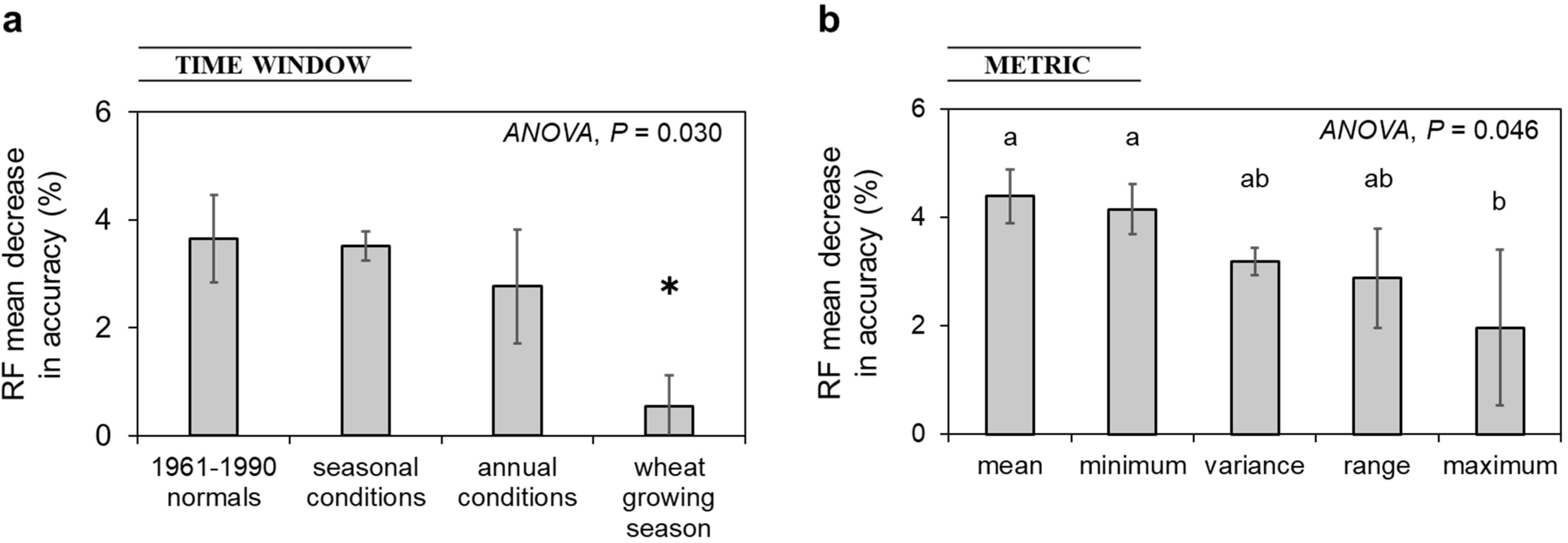
Characterisation of the thermal niche at each sampling site. The mesoclimatic thermal environments of the 8 Euro-Mediterranean sampling locations were summarised (a) for four climatic time windows (‘*1961-1990 climate normals*’ and thermal conditions for the year of sampling averaged over the calendar year *i.e.* ‘*annual conditions*’, the ‘*wheat growing season*’ *i.e.* from late October to early July, the ‘*seasonal conditions*’ *i.e.* contrasts between the winter and spring periods) and (b) with five metrics (thermal mean, range, maximum, minimum and variance). The 20 thermal variables were ranked according to their contributions to overall variations in temperature between the three contrasting Köppen-Geiger climatic zones prospected. For the establishment of a thermal niche classification, the most discriminating variables were identified by applying a nonlinear and nonparametric random forest algorithm (RF; Breiman, 2001) with the ‘*randomForest*’ package of R (n_tree_ bootstrap samples = 1000; Liaw & Wiener, 2002). The importance of the variables was compared based on the RF mean decrease in accuracy, which provides a metric indicating the extent to which the exclusion or permutation of a variable reduces RF accuracy (classification error). Higher mean decreases in accuracy for a given variable indicate a greater importance of the variable concerned for the classification of climatic zones. One-way analyses of variance (ANOVA outputs displayed with letters indicating the results of Tukey post-hoc multiple pairwise-comparisons) indicate that (a) the climatic time window of the wheat growing season and (b) maximum temperature are significantly less informative for classifying the three climatic zones.

**Table S1.**
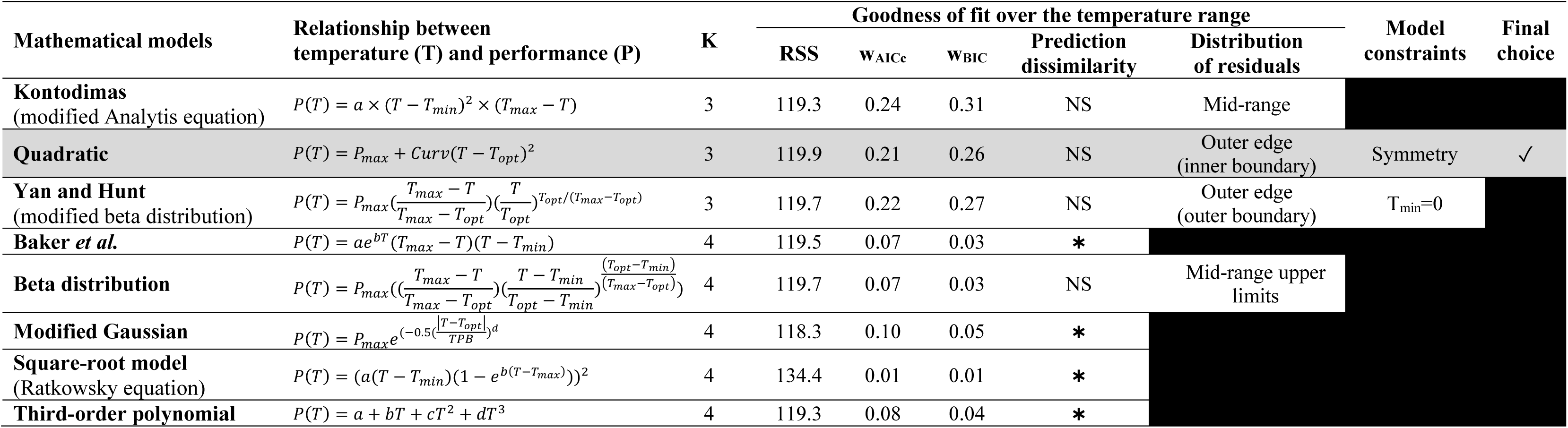
Selection of a candidate mathematical model for establishing TPCs. Model equations relating performance (P) to temperature (T) by means of K parameters were harmonised with the following key thermal parameters, when relevant: P_max_ (maximum performance), T_opt_ (thermal optimum), T_min_ (minimum temperature), T_max_ (maximum temperature), *Curv* (shape parameter) and TPB (thermal performance breadth). As in Boixel *et al*. (2019), the most appropriate model was selected to fit the TPC of each strain on the basis of: (i) statistic metrics accounting for goodness-of-fit namely residual sum of squares (RSS), Akaike weight (w_AIC_) and Schwarz weight (w_BIC_); (ii) the similarity between equation predictions of thermal responses over the temperature range and the best-fitting model (prediction dissimilarity: NS: not significant; ✱: *P* < 0.05) and their corresponding distribution of residuals along the temperature axis (with a view to maximising the accuracy of estimation for T_opt_ and the surrounding supra- and suboptimal estimates over the thermal extremes T_min_ and T_max_); (iii) model constraints potentially forcing the fitting process in a particular direction. Successive steps in the selection process are represented from the left to the right side of the table (black cells display discarded models). In this study of the thermal responses of *Z. tritici* over a Euro-Mediterranean spatio-temporal scale, as in the initial study performed in France, no significant difference in fit quality was observed between the quadratic and beta models for growth rate data over the entire range of temperatures investigated (from 6.5 to 33.5°C). However, their residuals were not similarly distributed along the temperature-axis, leading to small differences between equations in estimation accuracy for performance at mid-temperature ranges relative to outer edges (see the column ‘distribution of residuals’ below and in ESM2 of Boixel *et al*., 2019). In both studies, we selected the model that best estimated performance over the mid-temperature range (here the quadratic model), as diversity was assessed on three thermal traits extracted from this range of the TPC (P_max_, T_opt_, TPB_80_).

**Table S2.** Population-pairwise genetic distance (matrix of F_ST_ values). Pairwise comparisons between the 8 geographic *Z. tritici* populations (DK: Denmark; FR: France; IR: Ireland; IS: Israel; KZ: Kazakhstan; LV: Latvia; RU: Russia; TN: Tunisia) and 4 seasonal *Z. tritici* subpopulations (WIN1, WIN2: post-winter; SPR1, SPR2: post-spring). The matrix displays the estimation of pairwise F_ST_-values for each combination of populations based on 12 neutral microsatellite markers. Values in italics indicate significant F_ST_ values (*P* < 0.05), as evaluated with random allelic permutation procedures (1,023 permutations). These low population pairwise F_ST_-values and associated statistical analyses (conducted with the ARLEQUIN program; see Methods S3) indicate an absence of genetic differentiation between populations based on neutral markers (exact global test of sample differentiation based on haplotype frequencies conducted with 100,000 Markov steps: *P* = 0.75 ± 0.11)

|  | WIN2 | WIN1 | SPR2 | SPR1 | DK | FR | IR | IS | LV | KZ | RU | TN |
| --- | --- | --- | --- | --- | --- | --- | --- | --- | --- | --- | --- | --- |
| WIN2 | 0 |  |  |  |  |  |  |  |  |  |  |  |
| WIN1 | -0.013 | 0 |  |  |  |  |  |  |  |  |  |  |
| SPR2 | -0.007 | -0.008 | 0 |  |  |  |  |  |  |  |  |  |
| SPR1 | -0.012 | -0.014 | *-0.002* | 0 |  |  |  |  |  |  |  |  |
| DK | *-0.005* | *-0.005* | *0.001* | *0.000* | 0 |  |  |  |  |  |  |  |
| FR | -0.006 | *-0.004* | *-0.004* | *-0.005* | *-0.001* | 0 |  |  |  |  |  |  |
| IR | -0.007 | -0.006 | *-0.006* | *-0.002* | -0.006 | -0.006 | 0 |  |  |  |  |  |
| IS | *0.012* | *0.010* | *0.008* | *0.016* | *0.001* | *0.004* | *-0.000* | 0 |  |  |  |  |
| LV | *0.024* | *0.022* | *0.031* | *0.021* | *0.021* | *0.021* | *0.023* | *0.033* | 0 |  |  |  |
| KZ | -0.012 | -0.010 | *-0.002* | -0.009 | *-0.002* | *-0.004* | *-0.005* | *0.019* | *0.018* | 0 |  |  |
| RU | *0.004* | *0.006* | *0.009* | *0.005* | *0.010* | *0.006* | *0.008* | *0.024* | *-0.001* | *-0.001* | 0 |  |
| TN | *0.014* | *0.014* | *0.015* | *0.010* | *0.022* | *0.012* | *0.017* | *0.020* | *0.033* | *0.015* | *0.015* | 0 |

**Methods S1.**
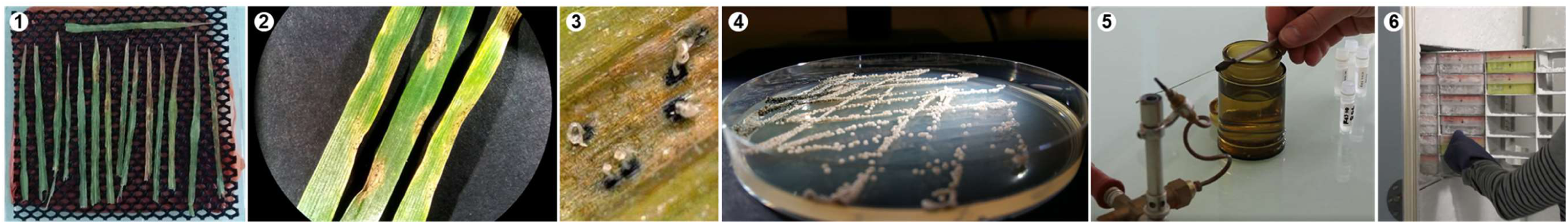
Procedure for the sampling, collection and recovery of *Zymoseptoria tritici* strains. Leaves with Septoria tritici blotch lesions were sampled at the various field locations, dried and flattened naturally for 2-3 days between two pieces of blotting paper, (1) placed overnight at 18 °C in a humidity chamber to promote cirrhus exudation and then (2) observed with a binocular magnifier to visualise pycnidia. (3) On each leaf, one cirrhus from a single pycnidium was retrieved from a randomly sampled single lesion, for further isolation in pure culture through (4) subculturing a single colony on PDA (potato dextrose agar, 39 g L^−1^) at 18°C in the dark. (5) After purification, *Z. tritici* spore suspensions were stored in stock tubes, in a 1:1 glycerol–water mixture, and were added to (6) the INRA BIOGER *Z. tritici* collection, which is stored at −80 °C.

**Methods S2.**
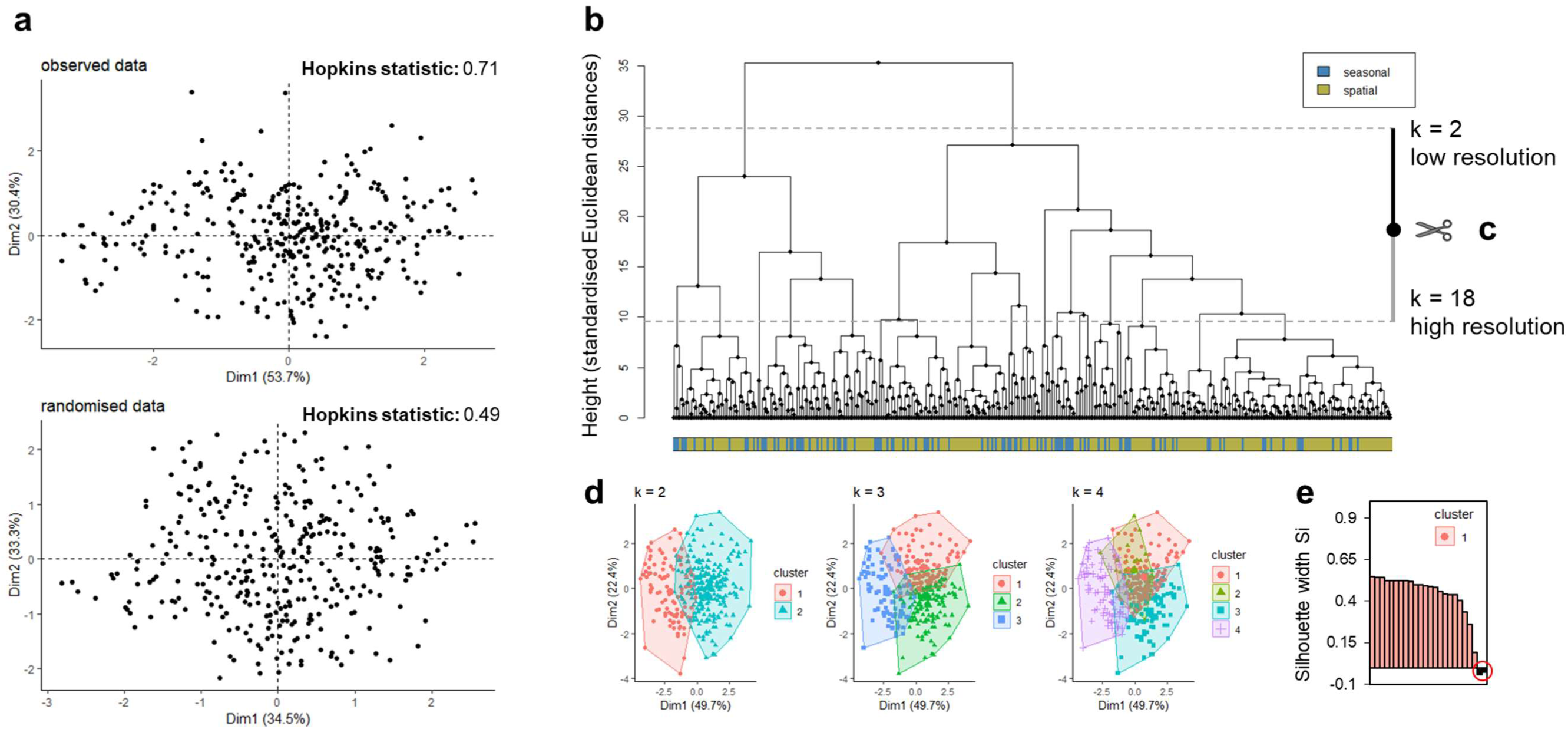
Definition of *Z. tritici* ‘thermotypes’. Thermotypes should be considered here as functional groups of thermal performance curves (TPCs) with similar thermal sensitivity features. Thermotype definition involves the establishment of a typology accounting for the diversity encountered in a given data set. This typology was built on the standardised phenotyping multidimensional data (including Pmax, Topt and TPB80 for both seasonal and geographic populations) in a five-step approach: (a) testing for uniformity in the data by calculating the Hopkins statistic (which measures spatial randomness and, thus, the tendency of a given data set to cluster; Lawson & Jurs, 1990) to determine whether a given data set can be divided into meaningful clusters; (b) calculating Euclidean distances between pairs of individual phenotypes and building the corresponding distance-based tree; (c) determining the optimal number of clusters (the cut-off being represented by the pair of scissors) by varying k from k = 2 (as we highlighted that there are clusters in the data set; see the Hopkins statistic in S2a) to k = 18 (to account for all combinations of variables: P_max_ × T_opt_ × TPB_80_). We achieved this by data mining, by identifying the best clustering scheme using 30 clustering validity indices implemented in the ‘*NbClust*’ R package (Charrad *et al*., 2014); (d) K-means clustering, an iterative algorithm with the objective function of minimising the pooled mean distances within clusters. This approach identified the 13 clusters to be generated into which the data could be partitioned (‘*kmeans*’ function with 13 centers and 50 initial configurations); (e) assessing the quality of the clustering result, by determining how well observations clustered with the ‘*silhouette*’ function of the ‘*cluster*’ package of R (Maechler *et al*., 2018). Based on this silhouette analysis (Rousseeuw, 1987), outliers among the thermal responses clustered (i.e. wrongly assigned phenotypes with a negative silhouette width coefficient – circled in red in the figure) were reassigned to the neighbouring cluster. This final partitioning of data (clustering result with reassigned outliers) was used to compare the corresponding clusters (*i.e.* thermotypes) on the basis of their features (compactness, separation, connectivity within and between clusters), abundance and distribution pattern within and between the 12 *Z. tritici* populations.

**Methods S3.**
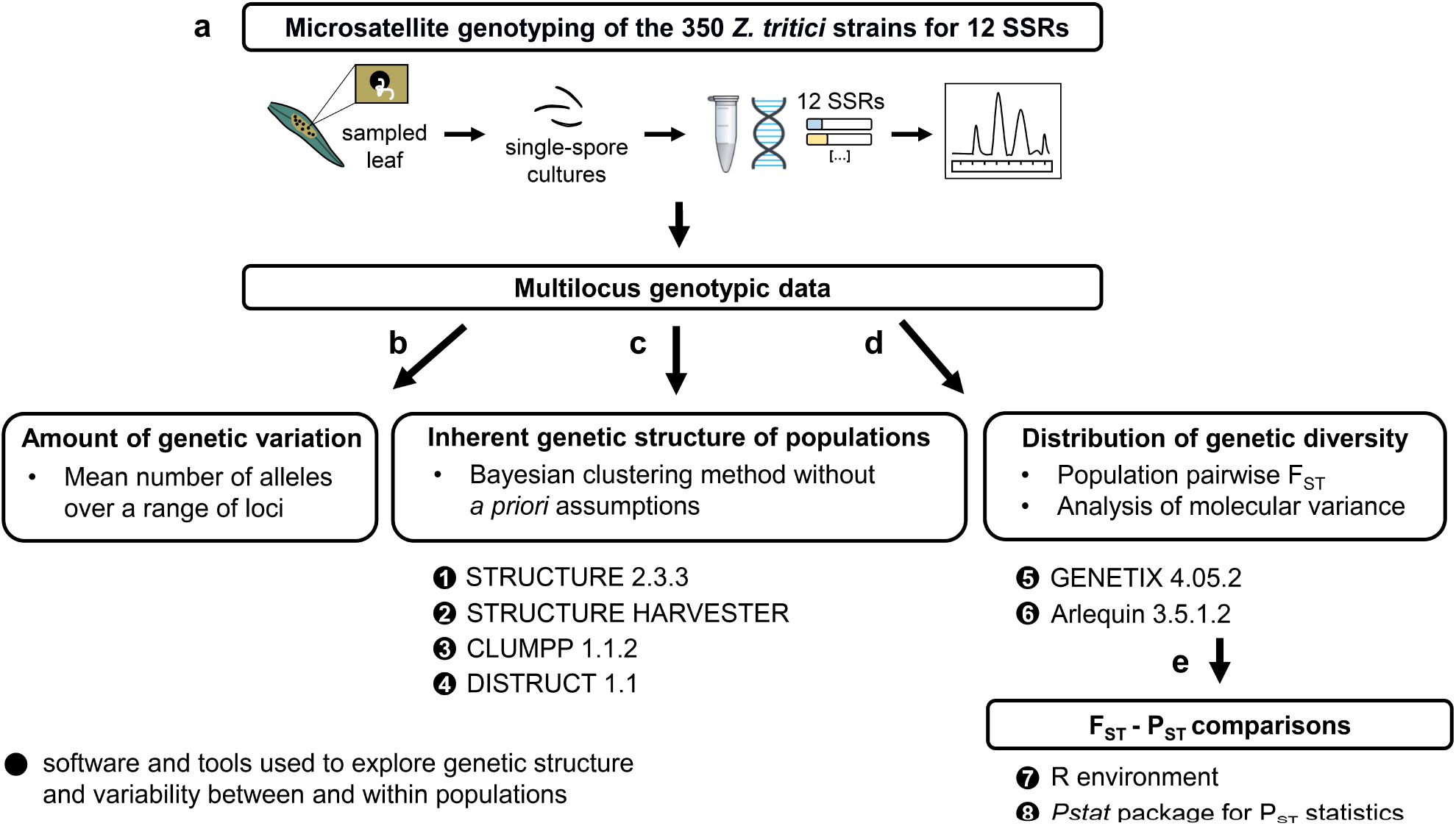
Procedure for acquiring and analysing multilocus genotypic data. (a) Genetic variability, structure and the distribution of diversity between and within the 12 *Z. tritici* populations were assessed based on microsatellite genotyping data. The overall data set was acquired for 12 SSRs (ST1, ST2, ST3A, ST3B, ST3C, ST4, ST5, ST6, ST7, ST9, ST10, ST12 neutral microsatellites; Gautier *et al*., 2014) following: (i) DNA extraction from single-spore cultures with the DNeasy Plant Maxi Kit (QIAGEN); (ii) amplification and sequencing of the SSR markers in one multiplex PCR sample (Eurofins Analytics France); (iii) the determination and annotation of allele sizes through visual analysis of individual chromatograms with Peak Scanner (Applied Biosystems). (b) Genetic variation in allelic distributions was assessed manually by the direct counting of the mean number of alleles (MNA) observed over all loci. (c) The distribution of this genetic variation within and between populations (population structure) was inferred with the Bayesian clustering approach implemented in STRUCTURE (Pritchard *et al*., 2000) under the admixture and correlated allele frequencies model. The algorithm was run on the basis of 500,000 iterations of the Markov chain (‘burn-in length’) followed by a run phase of 1,000,000 iterations (‘burn-in period’) with 10 independent replicates for each tested number of clusters (range set from 1 to 10). The optimal number of inferred genetic clusters (K) was estimated by the Evanno method (Evanno *et al*., 2005) with STRUCTURE HARVESTER (Earl & vonHoldt, 2012). Output data were then visualised with CLUMPP (Jakobsson & Rosenberg, 2007) and DISTRUCT (Rosenberg, 2003). (d) The genetic divergence between the sampled populations (genetic differentiation assessed on pairwise estimates of Weir and Cockerham’s F-statistic: 1,000 randomisations) was calculated with GENETIX (Belkhir, 2004). Hierarchical analyses of molecular variance (AMOVA) were conducted with Arlequin (Excoffier & Lischer, 2010) to assess the contribution of sampling date and location to the differences and patterns of genetic variance detected. (e) For inferences of the contribution of genetic drift and natural selection to the variation in thermal responses between populations, P_ST_ values (phenotypic differentiation between populations) of P_max_ (maximum performance), T_opt_ (thermal optimum), TPB_80_ (thermal performance breadth) and their confidence intervals were calculated with the ‘*Pstat*’ package of R (using the ‘*TracePst*’ function under the arguments boot = 1000 and pe = 0.95; Da Silva & Da Silva, 2018). F_ST_-P_ST_ comparisons were performed to test for patterns of local adaptation and to assess whether phenotypic differentiation between populations were greater or smaller than expected under the influence of genetic drift.

